# Systematic quantification of synapses in primary neuronal culture

**DOI:** 10.1101/2020.02.17.952242

**Authors:** Peter Verstraelen, Gerardo Garcia, Marlies Verschuuren, Bob Asselbergh, Rony Nuydens, Peter Larsen, Jean-Pierre Timmermans, Winnok H. De Vos

**Author notes:** equal contribution (Peter Verstraelen, Gerardo Garcia). Correspondence: Prof. Dr. Winnok H. De Vos.

## Abstract

A vast set of neurological disorders is associated with impaired synaptic connectivity. Therefore, modulation of synapse formation could have therapeutic relevance. However, the high density and small size of synapses make their quantification a challenging task. To improve the reliability of synapse-oriented drug screens, we evaluated a panel of synapse-targeting antibodies for their labeling specificity on hippocampal and cortical cell cultures using quantitative immunofluorescence microscopy. For those antibodies that passed multiparametric validation, we assessed pairwise colocalization, an often-used readout for established synapses. We found that even when two pan-synaptic markers were used, the overlap was incomplete, and the presence of spurious signals limited the dynamic range. To circumvent this problem, we implemented a proximity ligation-based approach, that only leads to a signal when two pre- and postsynaptic markers are sufficiently close. We demonstrate that this approach can be applied to different synaptic marker combinations and can be successfully used for quantification of synapse density in cultures of different maturity stage in healthy or pathological conditions. Thus, the unbiased analysis of synapse labeling and exploitation of resident protein proximity, allows increasing the sensitivity of synapse quantifications in neuronal culture and therefore represents a valuable extension of the analytical toolset for *in vitro* synapse screens.

## Introduction

Synapses are the prime mediators of neuronal communication and their plasticity defines learning and memory. They are considered to be vulnerable to many neuropathological conditions. In Alzheimer’s disease for instance, soluble oligomers of beta-amyloid (Aβ) and hyperphosphorylated tau localize to synaptic terminals, causing their number to decline even in the early stages of the disease (1–3). Synapse loss is recapitulated in mouse models that overexpress human APP (4, 5) and in WT mice that have received an intracerebroventricular injection of Aβ oligomers (6). Since synapse loss is considered to adversely affect cognition, reversal or prevention thereof may represent a therapeutic strategy for neurodegenerative diseases (7).

Discovery of novel compounds that modulate synapse density most often initiates with *in vitro* screens. In its simplest form, screening is done on primary neuronal cultures, stained for a specific synapse marker (8–10). Among the applications, this approach has been used to document the synaptotoxic effect of Aβ_1-42_, and its prevention by an oligomerization inhibitor (10). However, using a single marker to identify synapses is complicated by the presence of non-synaptic (*e.g.,* vesicular) or degenerate synaptic structures. To focus more specifically on mature synapses, the colocalization between a pre- and postsynaptic marker has been introduced as readout (11–16). Several gene silencing screens have made use of this approach to identify regulators of excitatory and inhibitory synapses (12, 15). Yet, also this approach is not free from caveats, as the use of antibodies limits the sensitivity of the assay to their specificity. While elegant genetic strategies for synapse labeling have been conceived (17), from a screening perspective, immunostaining is preferred for its ease-of-use, universal applicability and lack of overexpression artefacts. One way to validate antibody specificity consists of knocking down the target gene (13, 15). However, such an approach is expensive and may not provide a sufficiently reliable result as the knockdown could be incomplete, off-target and alter the cellular phenotype (*e.g.,* cytotoxic). Hence, a more systematic validation of antibody performance that does not require experimental interventions would be welcome. To satisfy this need, we introduced a segmentation-independent microscopy image analysis of 28 commercially available antibodies raised against pre- and postsynaptic proteins on primary cortical and hippocampal cultures. We also quantified the degree of colocalization between pre- and postsynaptic markers and found significant differences in synapse count between different marker combinations. Finally, to improve the sensitivity and reproducibility of synapse density quantification, we introduced a transsynaptic proximity-ligation assay (PLA). Considering the unique property of PLA to only detect protein interactions at distances below 40 nm (18), we identified bonafide partners at the pre- and postsynaptic side. We show that transsynaptic PLA has the specificity and sensitivity necessary to provide reliable synapse counts for *in vitro* experiments aimed at synaptic modulation.

## Materials and Methods

### Primary neuronal cell culture

This study was carried out in accordance with the recommendations of the ethical committee for animal experimentation of the University of Antwerp (approved ethical file 2015-54). Hippocampi and cortex were dissected from WT C57Bl6 and PSD95-CreNABLED (19) (purchased from the Jackson laboratories, ref 029242). E18 mouse embryos in HEPES (7 mM)-buffered Hanks Balanced Salt Solution (HBSS-HEPES), followed by trypsin digestion (0.05%; 10 min; 37°C) and mechanical dissociation. After centrifugation (5 min at 200g), the cell pellet was resuspended in Minimal Essential Medium supplemented with 10% heat-inactivated normal horse serum and 30 mM glucose. Cells were plated in Poly-D-Lysin-coated 96-well plates (Greiner µClear), at 30,000 cells/cm^2^ (for immunocytochemistry), or in 6-well plates at 60,000 cells/cm^2^ (for western blot) and kept in a humidified CO_2_ incubator (37°C; 5% CO_2_). After 4 hours, the medium was replaced with 150 µl B27-supplemented Neurobasal medium (NB-B27), containing Sodium Pyruvate (1 mM), Glutamax (2 mM), glucose (30 mM) and Penicillin-Streptomycin (0.5%). To suppress proliferation of non-neuronal cells, 0.5 µM arabinosylcytosine was added in 25 µl NB-B27 at the third and tenth day *in vitro* (DIV). Cell culture supplies were purchased from ThermoFisher (Waltham, MA, USA).

### Western blotting

14 DIV cortical and hippocampal cultures were lysed using ice-cold RIPA buffer supplemented with phosphatase and protease inhibitor (HALT cocktail, ThermoFisher 78445) and 5 mM EDTA. The lysate was centrifuged (10.000g, 20 min, 4 °C) and the protein concentration of the supernatant was determined using a BCA assay (ThermoFisher 23225). Samples were denatured (70% sample, 25% LDS, 5% DTT) for 10 min at 70 °C before being loaded on a 4-12% Bis-tris gel (ThermoFisher NP0322BOX) at 10 µg/lane. A stained ruler was included in the first and last well (ThermoFisher 26616). The gel tank was filled with NuPage MOPS SDS running buffer and NuPage anti-oxidant, and was cooled during electrophoresis (200V, ± 1h). Proteins were transferred to a PVDF membrane using NuPage transfer buffer (30V, 1h). To check the transfer, blots were reversibly stained with a 0.1% Ponceau S solution in 5% acetic acid. Blots were subsequently blocked with 5% ECL blocking solution in Tris-buffered Saline with 0.1% Tween (TBS-T). Blots were cut into 5 pieces so that 2 lanes (hippocampal and cortical) with or without the ruler could be stained in 1 reaction. Primary antibody was applied overnight at 4 °C on a roller, followed by a TBS-T wash (3 x 5 min). Horse Radish Peroxidase-coupled secondary antibodies were incubated for 2h at room temperature (RT), followed by a final TBS-T wash. All antibodies were diluted in blocking buffer and are listed in **Suppl. Table 1**. After reconstructing the cut blots into their original positions, bioluminescent detection was performed using Immobilon Western HRP substrate (Merck Millipore WBKLS0500, 30 sec) and a Chemidoc Touch imager (Biorad, Temse, Belgium). After completion, the blots were restained for GAPDH as loading control. The global contrast of the individual (cut) blots was adjusted with Fiji image analysis freeware (20).

### Immunofluorescence staining (IF)

Paraformaldehyde-fixed cultures (2%, 20 min, RT) were permeabilized with 1% Triton X-100 in blocking buffer (0.1% bovine serum albumin and 10% normal horse serum in PBS) for 10 min, followed by an overnight incubation with the primary antibodies (**Suppl. Table 1**) at 4 °C in blocking buffer. After washing with PBS, secondary antibodies (**Suppl. Table 1**) were added for 2 hours. Finally, 4’,6-diamidino-2-phenylindole (DAPI) was applied to the cultures for 10 min at a concentration of 2.5 µg/ml, followed by a PBS wash. Primary antibodies targeting extracellular epitopes were incubated prior to permeabilization to reduce non-specific intracellular background staining. This was followed by a PBS wash and permeabilization to continue with antibodies targeting intracellular epitopes. All presynaptic markers were designated to the 561 nm excitation channel, while postsynaptic markers were labeled for the 488 nm channel. Secondary antibodies were kept identical where possible, depending on the species of the primary antibody.

### Expansion microscopy

The protocol for expansion of the samples was adapted from (21). In brief, immunostained samples were crosslinked for 10 min in 0.25% glutaraldehyde in PBS. Gelation was done in a mixture of 2M NaCl, 2.5% (w/w) acrylamide, 0.15% (w/w) N,N’-methylenebisacrylamide, 8.625% (w/w) sodium acrylate in PBS with polymerization initiated with TEMED and APS. Polymerized gels were incubated for 30 min at 37°C in a digestion buffer containing 8 U/ml proteinase K. Cover glasses were removed from the digested gels, which were placed in high volumes (> 30 mL) of distilled water that were exchanged at least 5 times until full expansion of the gels. The lateral expansion factor was determined by measuring the gel dimensions before and after expansion. Finally, the gels were trimmed, nuclei counterstained with Hoechst 33342 (1/5000 in water for 30 min at room temperature), positioned in 50 mm diameter glass bottom dishes (WillCo Wells GWSt-5040) and immobilized using 2% agarose.

### Dendritic spine labeling

To label spines in a sparsely distributed subset of neurons, 14 DIV cultures were incubated with the lipophilic dye CM-DiI (ThermoFisher C7000, 5 µg/ml, 20 min), followed by a PBS wash. The next day, after the dye had spread throughout the plasma membrane, the cultures were gently permeabilized using a glycerol gradient (50% glycerol for 20 min, 80% for 20 min and 100% for 50 min) followed by an extensive PBS wash. This permeabilization protocol preserved DiI fluorescence while allowing antibody penetration. A similar, but longer protocol was used for IF, in which the primary antibody was applied overnight at room temperature and the secondary for 4 hours.

### Synaptic Proximity Ligation Assay (synaptic PLA)

Primary cultures fixed and permeabilised as described for IF were used to perform PLA using a commercial kit (Duolink® PLA starter kit, Sigma-Aldrich) according to manufacturer’s instructions. Briefly, cells were incubated with a commercial blocking buffer, and then incubated overnight at 4°C, with primary antibodies diluted in the commercial solution at concentrations described in **Suppl. Table 1**. After PLA probe incubation, ligation and amplification steps, samples were immunostained for MAP2 and Synapsin for triple staining experiments, counterstained with Duolink® Nuclear Stain and mounted with anti-fade buffer (Sigma-Aldrich). Images were taken no more than 2 days after the PLA procedure to ascertain signal preservation.

### Microscopy

Cultures were imaged with a spinning disk confocal high-content imager (Opera Phenix, PerkinElmer) equipped with a robotic arm for plate loading. A 40X water immersion objective (numerical aperture (NA) 1.1) was used. At 488 nm excitation, the optical resolution of the system is 0.271 μm (and corresponding pixel size 0.149 μm), which is considerably larger than the distance of the synaptic cleft (15–25 nm), yet sufficiently small to allow signals of corresponding pre- and postsynaptic markers to partially overlap. Per well, 15 fields were acquired in 4 channels (405nm, 488nm, 561nm and 640nm excitation) at 4 axial positions separated by 1 µm spacing. Different fluorescence channels were separated using standard excitation/emission filters and dichroic mirrors. Owing to the large dynamic range of the Opera Phenix system, acquisition settings could be kept identical for all experiments. For assessing localization of postsynaptic markers in dendritic spines and synaptic PLA triple stainings (synaptic PLA/mVenus/synapsin), a spinning disk confocal research microscope (UltraVIEW VoX, PerkinElmer) was used, equipped with a 60X water objective (NA 1.2) resulting in an optical resolution of ∼200 nm and corresponding pixel size of 120 nm. Finally, for super-resolution radial fluctuations (SRRF) microscopy (22), the same setup was used to acquire stacks of 200 images of the same confocal plane, using the Perfect Focus System to prevent focus drift.

### Image analysis

To assess the degree of colocalization between two synaptic markers, we adopted the *van Steensel* approach (23). This method calculates the Pearson correlation coefficient (PCC) between two images as a function of a lateral shift (Δx), resulting in a correlation function (CF). When the image is compared to its (shifted) duplicate, the result is an auto-correlation function (ACF). In case of complete colocalization, the correlation function (CF) has a maximum (PCC = 1) at Δx = 0 (no shift); in case of exclusion, it reaches a minimum at this position. The shape of the CF not only informs on the presence or absence of colocalization, when colocalization is present, it also provides information on the size of the overlapping spots (the width of the CF measured as full width at half maximum (FWHM)) and the dynamic range – and thus quality – of the staining (measured as the amplitude of the CF). Exactly because of this latter property, the approach was also used to assess the specificity of single synapse marker staining, the assumption being that the ACF of an image decreases with Δx much stronger for a specific staining than it does for less specific ones. The maximal pixel shift used for both analyses was 50 pixels in both directions, which is well beyond the typical spot diameter of ∼ 10 pixels. The variation across images, or the spread, was expressed by a band of 1 standard deviation above and below the average CCF and measured at dx = 50 px for plotting. All measurements were performed in FIJI image processing freeware using a home-written script that is available upon request.

For spot and synapse density quantification, multidimensional images were analyzed using a previously developed image analysis pipeline for Acapella software (PerkinElmer) (16). In brief, nuclei (DAPI) and neurites (MAP2) were segmented based on a user-defined fixed threshold. Non-neuronal nuclei were removed based on their larger size, lower circularity and limited overlap with the neurite mask. Next, both the neurite and neuronal nuclei mask were dilated and subtracted from each other to obtain a search region for synapse marker spots. Spots were enhanced using a Difference of Gaussian filter with a ratio of 1.6. To set thresholds in an unbiased manner, a Fiji script was written that allowed user-friendly interaction with the images while being blinded for the synapse marker at hand. Spot counts were normalized to the neurite area to obtain the spot density. For double stainings and PSD-mVenus experiments, the density of colocalizing spots in 2 fluorescence channels was calculated as those spots for which at least 1 pixel overlaps. We previously determined that measurements were not biased by chromatic aberration or the overlap criterion (16). For colocalization of antibodies with PSD95-mVenus, the percentage of PSD95-mVenus spots that have an overlapping synapse marker spot was measured. For double immunostainings, the percentage of all spots that reside in synapses was calculated as the density of colocalizing spots divided by the density of all (colocalizing and non-colocalizing) spots multiplied by 100.

For double labeling with dendritic spines, the puncta within a CM-DiI-positive stretch were counted manually on anonymized images and assigned to either the dendritic shaft or spine. The percentage of spots residing in dendritic spines, and the percentage of spines containing spots were calculated. For quantitative distance analysis in synaptic PLA images, center coordinates were extracted for all spots, and the pairwise distance distribution between spot types (PLA, mVenus, Synapsin) was analyzed in R using the package “spatstat” by applying the nearest neighbor correlation (nncorr) method after correcting for the search region (i.e., the neurite mask) (24). For SRRF, the NanoJ plugin for FIJI was used (22). Briefly, after drift correction with NanoCore, a temporal radial auto-correlation of order 2 was applied, with gradient smoothing, intensity weighting and gradient weighting (PSF FWHM = 3.17) for spot channels, and the previous settings with renormalization for the MAP2/DAPI channel.

### Data visualization and statistics

Graphs were constructed in Graphpad Prism and statistics performed in SAS JMP. For single, double and ENABLED stainings, 6 wells were considered per condition (marker or marker combination and cell type). Within each well, 15 fields were imaged. For statistical comparison of spot densities, these fields were averaged so that 1 data point represented 1 well. A one-way ANOVA was performed using marker as fixed factor followed by post-hoc testing for predefined comparisons (antibodies that had the same target; 7 comparisons) with Sidak’s correction. For triple stainings and hTau-P301L overexpression experiments, two-way ANOVA was performed, with marker and PLA type or MOI and method (PLA or colocalization) as fixed factors, respectively. Van Steensel analyses were performed on individual fields, amounting to 90 data points per condition, while segmentation results were averaged per well before plotting (6 data points per condition). For dendritic spine analysis, at least 200 spines were counted per condition. For synaptic PLA experiments a minimum of 4 wells with 15 fields per well were averaged per condition. For SRRF of synaptic PLA staining, between of 4 and 7 images were used per combination.

## Results

### Not all synapse markers can be labeled with equal specificity

Synapses harbor a unique proteome for which a wealth of dedicated antibodies has been developed. Yet, despite their ample use, the specificity and generalizability of many remains questionable. Therefore, we screened a panel of synapse-targeting antibodies for their labeling specificity on hippocampal and cortical cultures at 14 days *in vitro* (DIV), a time point at which synaptic connections are well-established (16, 25–27). Targets included proteins of pre- (vesicular, active zone) and postsynaptic (receptor, scaffold) compartments, as well as trans-synaptic adhesion proteins (**Suppl. Table 1, Fig. 1a, b**). When quantifying the absolute spot density using a user-defined threshold, we found variable results in that antibodies targeting the same antigen yielded significantly different spot densities (Neurexin, Neuroligin1, GluN1) or a very large variability between replicates (PSD95 (a), Shank) (**Fig. 1a** for cortical and **Suppl. Fig. 1** for hippocampal cultures). Exemplary, the sum of excitatory (vGLUT) and inhibitory (vGAT) spots was not equal to the number of spots stained by a pan-presynaptic marker (Synapsin, Synaptophysin, Bassoon). This suggests that not all antibodies stain with equal specificity and that spurious signals may represent a serious confounding factor.

**Figure 1.**
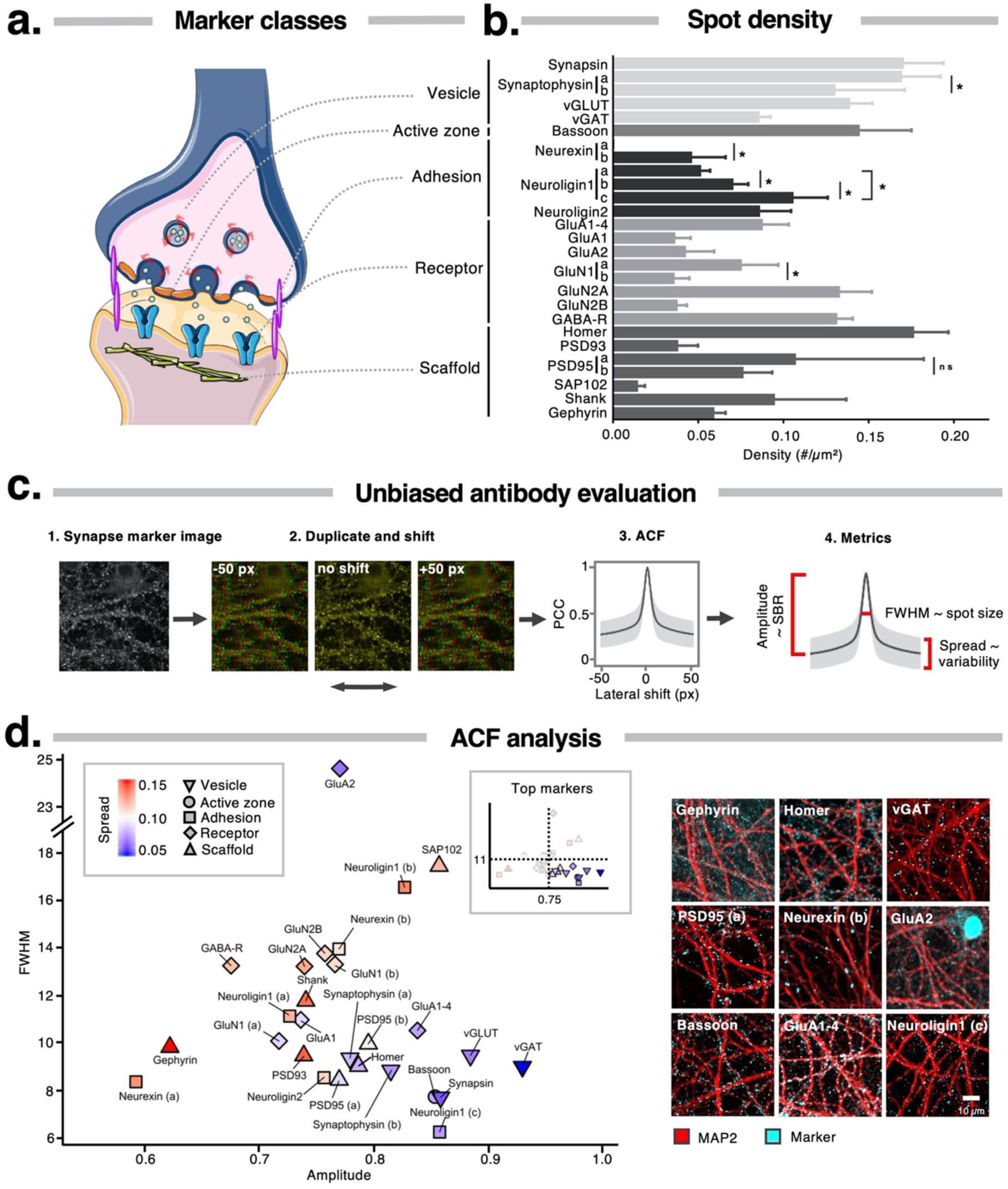
Antibodies targeting synapse markers have variable specificity. **a.** Markers of different functional classes considered in this work; **b.** Quantification of the spot density after immunostaining yields variable results even between antibodies that target the same marker (* p < 0.05, one-way ANOVA post-hoc Sidak’s multiple comparisons test); **c.** The auto-correlation function (ACF) allows unbiased evaluation of staining performance, by calculating the Pearson’s Correlation Coefficient (PCC) between an image and its duplicate as a function of a lateral shift. The ACF amplitude and spread report on the signal-to-background ratio and the variability across images, respectively, the full width at half maximum (FWHM) correlates with spot size. A crisp synapse spot image has a high amplitude (> 0.75) with narrow spread (< 0.08) and small FWHM (≤ 11), whereas a noisy image has a small amplitude with large standard deviation and large FWHM; **d.** Scatterplot of ACF parameters for the different synapse antibodies tested on primary cortical cultures at 14 DIV, along with representative images for a selected subset. Inset shows the same scatterplot after gating for optimal parameters.

To rule out bias arising from user-dependent thresholding, we adopted a method for unbiased analysis of IF staining performance, by determining the auto-correlation function (ACF) using the van Steensel approach (23) (see M&M; **Fig. 1c**). The amplitude of the ACF was used as proxy for labeling specificity, the full width at half max (FWHM) reported on the average spot size, whereas the spread (2 x standard deviation at dx = 50 px) around the average ACF informed on the variability between replicates. Based on the dimensions and quality of representative markers, we determined a window in which antibodies were considered to yield an optimal staining result (ACF amplitude > 0.75, FWHM < 11 (= expected spot size) and spread <0.08). All synaptic vesicle and active zone markers met these criteria (**Fig. 1d** for cortical and **Suppl. Fig. 1b** for hippocampal cultures, see **Suppl. Fig. 2** for the actual ACF plots). Except for Neuroligin1 (c), GluA1-4, Homer and PSD95 (a,b), antibodies that target adhesion molecules, neurotransmitter receptors and scaffold proteins yielded more blunted (amplitude < 0.75; FWHM > 11) and variable (spread > 0.08) ACFs, indicative of lower labeling specificity (**Fig. 1d**). Antibodies that target GluA2 and SAP102 yielded exceptionally high FWHM since they also labeled nuclei. For some inhibitory markers (Gephyrin and Neuroligin2), we observed considerable differences in the ACF of hippocampal and cortical cultures, which could point to variations in antigen abundance and inhibitory contacts between the two cell culture types (**Fig. 1d, Suppl. Fig. 1b**).

To gain more insight into the specificity of the same synapse antibodies, we performed western blots on cell extracts of 14 DIV cortical and hippocampal cultures (**Suppl. Fig. 3**). Only a minority (4/27; Neurexin (a), Neuroligin1 (a and b), and Shank) of the immunoblot-compatible antibodies failed to label a specific band near the predicted molecular weight of the targeted marker. Neurexin (a) antibody also had a suboptimal ACF, underlining its complete lack of specificity. However, the ACF of Neuroligin1 (a) and (b) and Shank did not show major deviations, suggesting that epitopes may have been lost upon denaturation for western blots. For several markers, multiple bands were commonly detected. These observations need not necessarily imply non-specificity as it may be the result of the extraction/denaturation procedure (exposing masked epitopes) or the presence of alternative isoforms. However, recognition of different isoforms may contribute to variable staining specificity. Notable examples include a Synapsin1/2 antibody which revealed many more isoforms in hippocampal than in cortical culture, and a vGAT antibody, which not only binds the full-length (synaptic) protein, but also a truncated, extra-synaptic variant (28). We conclude from these experiments that immunoblotting does not contribute significantly to the understanding of how antibodies will perform in immunofluorescence, as opposed to unbiased ACF analysis.

### Genetic and spine labeling strategies reveal partial synaptic localization

While IF showed that most antibodies yield a punctate staining, it was not clear whether the obtained signals solely represent synapses. That is why we sought for a strategy that could provide more certainty and serve as benchmark for determining synaptic localization. As a first approach, we made use of PSD95-mVenus (ENABLED) transgenic mouse cultures (19). In this model, endogenous PSD95 should be labeled with minimal overexpression artefacts. As we noticed the PSD95-mVenus signal increased with culture age (**Suppl. Fig. 4**), we analyzed the colocalization at both 14 and 28 DIV (**Fig. 2b, Suppl. Fig. 5-7**), using the van Steensel approach. In this case the CCF amplitude reports on the degree of colocalization and the FWHM on the combined size of the spots, as evidenced by simulations (**Fig. 2a**). We also measured the percentage of PSD95-mVenus spots that have an overlapping synapse marker spot using a segmentation-based approach (defined as overlap coefficient; OC). To determine the maximum level of colocalization attainable with both approaches (CCF and segmentation-based), an anti-GFP antibody (which cross-reacts with mVenus) was included as positive control. This control yielded a CCF amplitude of ∼ 0.7 and FWHM ∼ 10, and upon segmentation, an OC of ∼ 80% in cortical cultures at both DIVs (**Fig. 2b,** see **Suppl. Fig. 5** for hippocampal). Given that PSD95-mVenus is an excitatory marker, the amplitude of the CCF and OC were significantly larger for excitatory (*e.g.,* Homer) than for inhibitory (*e.g.,* vGAT, GABA-R) markers at both DIV. Surprisingly, IF with two validated PSD95 (a,b) antibodies – considered positive control as well – only resulted in modest colocalization with the PSD95-mVenus spots. Also, the synaptic adhesion proteins Neurexin (b), Neuroligin1 (c) and Neuroligin2 showed low colocalization with PSD95-mVenus. Of the tested presynaptic markers, Bassoon yielded the highest colocalization with PSD95-mVenus, consistent with its presence in the active zone, whereas vesicle markers (Synapsin, Synaptophysin (a), vGLUT) were further away from the PSD95-mVenus spots (resulting in higher FWHM) and therefore showed lower colocalization (lower amplitude and OC). This benchmarking indicated that the Homer antibody is best suited to label the excitatory postsynaptic compartment and that Bassoon is the preferred presynaptic marker.

**Figure 2.**
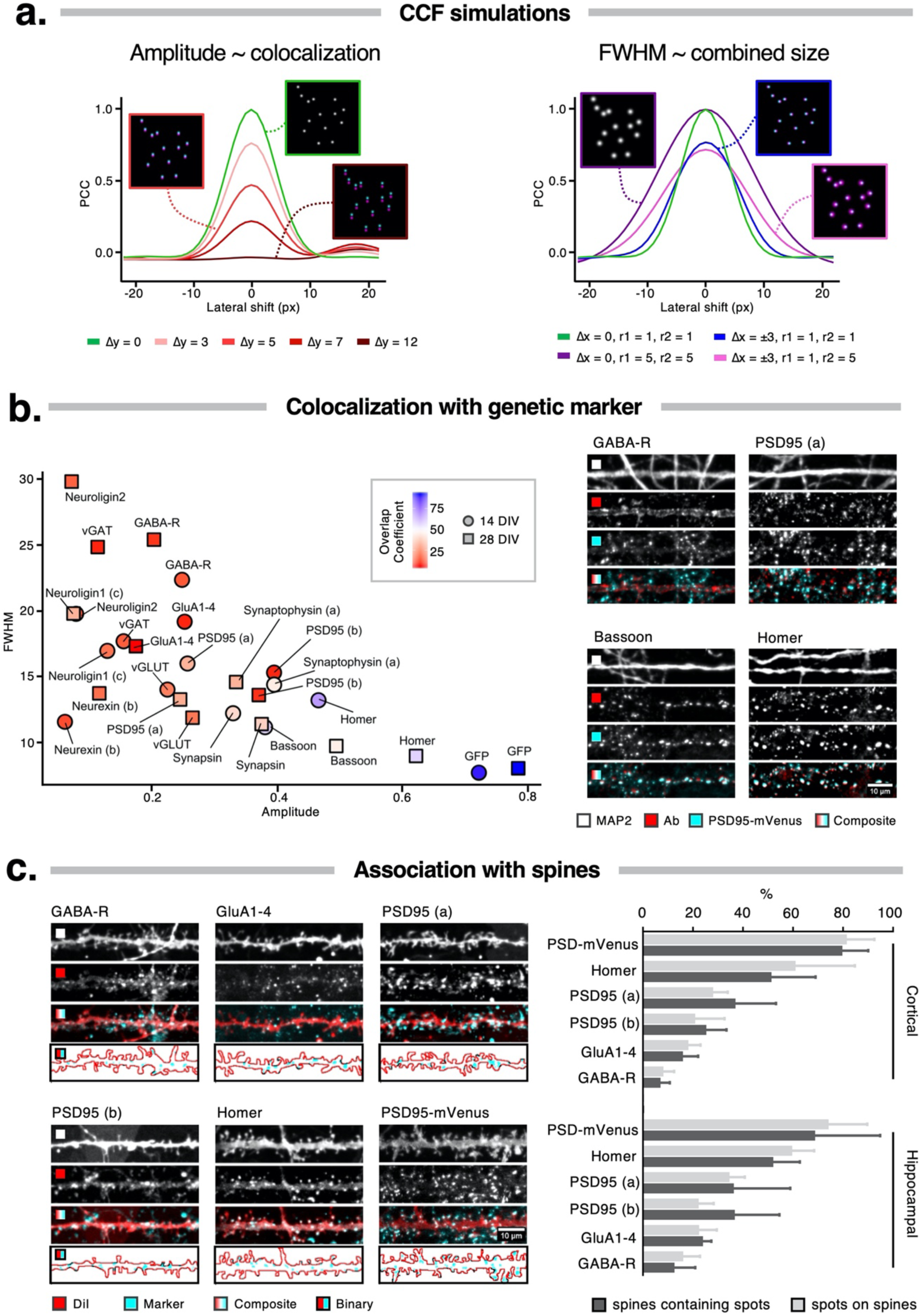
Association of synapse markers with the postsynaptic compartment. **a.** Cross-Correlation Functions (CCFs) for simulated spot images. An increasing mismatch between images (simulated by means of global lateral translation in y by 3, 5, 7 or 12 pixels) results in a progressively lower amplitude, while the FWHM increases with the size of the spots in at least one of both channels and is less sensitive to mismatch (simulated by randomly shifting spots 3 pixels up or down); **b.** Colocalization of synapse marker stainings with the genetic label PSD95-mVenus in 14 and 28 DIV primary cortical cultures, shown in a scatter plot from CCF parameters and color-coded by the overlap coefficient, defined as the percentage of PSD95-mVenus spots that have an overlapping signal from an antibody spot. An anti-GFP antibody, which cross-reacts with mVenus was used as a positive control. Representative images are from 28 DIV cortical cultures; **c.** Exemplary dendrite stretches from 14 DIV cortical cultures after co-labeling with postsynaptic synapse markers and DiI. Quantification of the association with spines was done in both cortical and hippocampal cultures.

In a second attempt to validate synaptic localization, we specifically focused on the excitatory postsynaptic compartment. This part of the synapse has a unique structure that is referred to as a dendritic spine (29). Despite significant morphological plasticity, spines can be readily visualized, among others using lipophilic dyes such as DiI (30, 31). When combining DiI with postsynaptic antibody staining, we found considerable differences in colocalization between synapse markers in cultures of 14 DIV (**Fig. 2c**). At the lower limit of the dynamic range, GABA-R puncta were rarely located on spines (6±4% for cortical and 12±9% for hippocampal), which corresponds to findings on cortical tissue at the ultrastructural level (32, 33). To gauge the upper limit of the dynamic range, we considered the genetic PSD95-mVenus signal and found that nearly 80% of the spots were located on spines, and that 80% of all spines contained mVenus signal. AMPA-R clusters, as labeled with GluA1-4 antibody, were partially located on dendritic spines, yet the majority of spots were found on the shaft. For PSD95 (a, b) antibodies, of which we theoretically expected spine labeling similar to PSD95-mVenus (∼80%), colocalization was less than half (30-40%) of the anticipated result. This implies that the antibodies did not label their target antigens with sufficient selectivity, as also suggested by the PSD95-mVenus colocalization measurements (**Fig. 3b**). With ∼50% of spines being labeled and ∼60% of all spots residing on spines, the scaffold protein Homer approached the upper limit of the dynamic range, proving more specific than both PSD95 antibodies.

**Figure 3.**
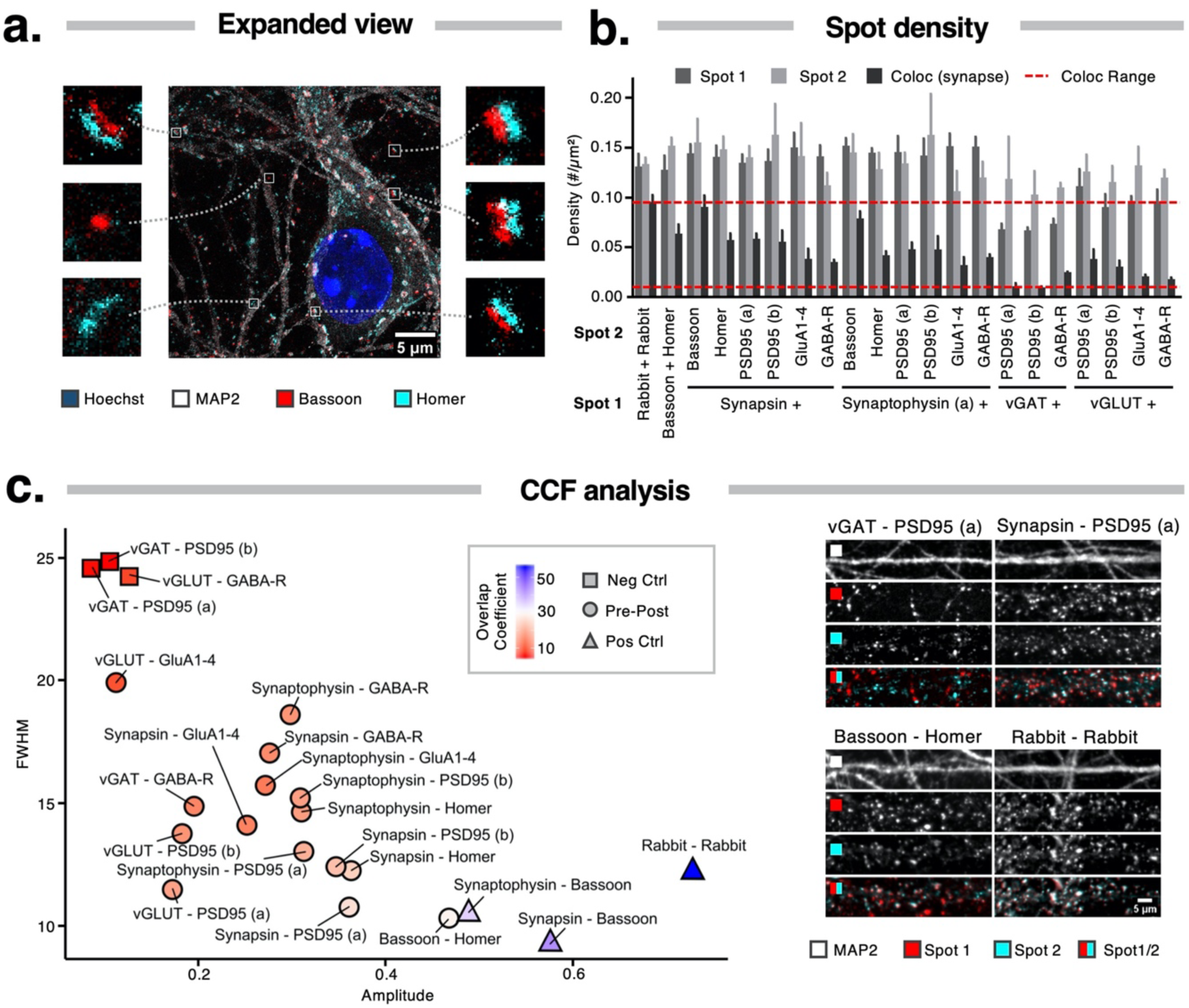
Quantification of synapse marker colocalization improves selectivity for mature synapses. **a.** Expansion microscopy shows the presence of juxtaposed pre- and postsynaptic signals in 14 DIV cortical cultures, as well as single marker spots; **b.** Quantification of spot and synapse density in 14 DIV cortical cultures shows the fraction of colocalized signals, assumed to represent mature synapses; **c.** Scatter plot of CCF parameters reporting on the colocalization of marker pairs in 14 DIV cortical neurons. The color code indicates the overlap coefficient, defined as the percentage of spots that reside in synapses. Pairs of inhibitory with excitatory synapse markers were considered negative controls while 2 pan-presynaptic markers and primary antibodies raised in the same species were used as positive controls.

Taken together, there is a substantial fraction of the IF signal that does not align with synapses, and, not all synapses are efficiently labeled by antibodies. Of those antibodies tested, Homer showed the highest colocalization with the PSD95-mVenus signal and strongest enrichment in dendritic spines, indicating that this antibody is best suited to stain the majority of the postsynaptic compartments.

### Synapse marker colocalization improves selectivity for mature synapses

The previous experiments revealed that synaptic antibodies label synapses with variable specificity and that non-synaptic spots are present, which complicates robust synapse quantification. Yet, using expansion microscopy, we confirmed that pre- and postsynaptic markers are often found juxtaposed in cultures at 14 DIV (**Fig. 3a**). The diffraction limit inherent to conventional fluorescence microscopy precludes signals closer than 200 nm from being resolved. With 20 nm, the synaptic cleft is well below this limit. Thence, a simple approach to detect mature synapses – *i.e.,* synapses that dispose of a clear pre- and postsynaptic compartment – with more certainty consists of quantifying the apparent overlap between pre- and postsynaptic markers (12, 13, 15, 16). We applied this method to synaptic marker antibodies that showed a crisp punctuate staining (evidenced by the ACF criteria defined above) and were raised in different species so that they could be combined in double stainings (**Fig. 3** for cortical, **Suppl. Fig. 8** for hippocampal cultures). Threshold-dependent spot quantification revealed that only a fraction of the pre- and postsynaptic spots resided in mature synapses, as evidenced by the lower density of colocalizing spots compared to the spot density of the individual markers (**Fig. 3b**, **Suppl. Fig. 8a**). The CCF between the two synapse marker channels was calculated, as well as the percentage of spots that reside in synapses (defined as the overlap coefficient, OC; **Fig. 3c**). The large FWHM (> 24 px), small amplitude (< 0.2) of the CFF and a low OC (< 5%) for combinations of inhibitory presynaptic and excitatory postsynaptic markers (vGAT and PSD95 (a/b)), or an excitatory presynaptic and inhibitory postsynaptic marker (vGLUT and GABA-R) illustrate that false positive detection by sheer chance is low (**Fig. 3c**, **Suppl. Fig. 8b**). As positive control, we used a combination of two rabbit polyclonal primary antibodies (vGAT and Homer), onto which the secondary antibodies were expected to bind with equal affinity. The resulting CCF had a large amplitude (> 0.65) and small FWHM (< 15 px; **Fig. 3c**, **Suppl. Fig. 8b**), while the CCF approached a value of 1 at Δx=0 (**Suppl. Fig. 9-10**), consistent with perfect colocalization. The OC on the other hand, amounted to only 56% for cortical and 44% for hippocampal cultures, again showing the strength of the unbiased CCF approach over spot segmentation. The colocalization (CCF and OC) was reduced when using two markers that label the same compartment (Synaptophysin (a)/Bassoon and Synapsin/Bassoon). Despite this limitation to the dynamic range, clear patterns could still be discriminated. For example, the combination of a pan-presynaptic marker (Synapsin or Synaptophysin (a)) with an inhibitory marker (GABA-R) could be distinguished from the combination with excitatory markers (PSD95 (a/b), Homer) (**Fig. 3c, Suppl. Fig. 8b**). The smaller FWHM, higher amplitude and larger OC indicate that the excitatory markers were closer to the presynaptic markers. Our results also indicated that GluA1-4 shows low colocalization with pan-(Synapsin, Synaptophysin) and excitatory (vGLUT) presynaptic markers, suggesting that many AMPA-R clusters reside outside of the synapse. In cortical as well as hippocampal cultures, Bassoon-Homer proved to be the combination with the highest colocalization, confirming that this combination is preferred for detecting mature synapses.

### Proximity of markers allows selective labeling of mature synapses

As the colocalization of two markers might be confounded by the presence of spurious signals (*e.g.,* extrasynaptic staining), we sought for an approach that would only give rise to positive signals in case pre- and postsynaptic markers are sufficiently close. We reasoned that the Proximity Ligation Assay (PLA) (34) would represent a plausible method to achieve this, as it only leads to a reaction when two markers (and their respective antibodies) are closer than 40 nm, a distance range that bridges the synaptic cleft (35) (**Fig. 4a**). We first validated the concept using a positive control – antibodies for lamin A and lamin B1, two nuclear envelope markers that are known to interact directly (36) – and a negative control – antibodies targeting lamin A and the mitochondrial marker TOMM20, which are spatially separated by much more than 40 nm. The positive control yielded nuclear-localized PLA spots, whereas the negative control produced virtually no signal, as was the case for a negative technical control in which only one of the primary antibodies was used (**Suppl. Fig. 11a**). Next, we tested whether targeting pairs of proteins present in excitatory and inhibitory synapses resulted in a synaptic PLA reaction. We found that the negative control - targeting vGLUT (excitatory) and Neuroligin 2 (inhibitory) – yielded a low PLA signal, whereas PLA for vGLUT and Neuroligin 1 (c) (both excitatory synapse markers) produced a strong punctate PLA pattern along the neurites, consistent with a synaptic staining. The synaptic PLA spot density fell within the range of values found for dual immunofluorescence against Homer/Bassoon (∼0.05 spots/µm^2^). We noted that the amount of synaptic PLA signal with the combination VGAT-Neuroligin1 (a discordant excitatory-inhibitory combination) was higher than for VGAT/Neuroligin2 which should detect reaction on inhibitory synapses only (**Suppl. Fig. 11b**).

**Figure 4.**
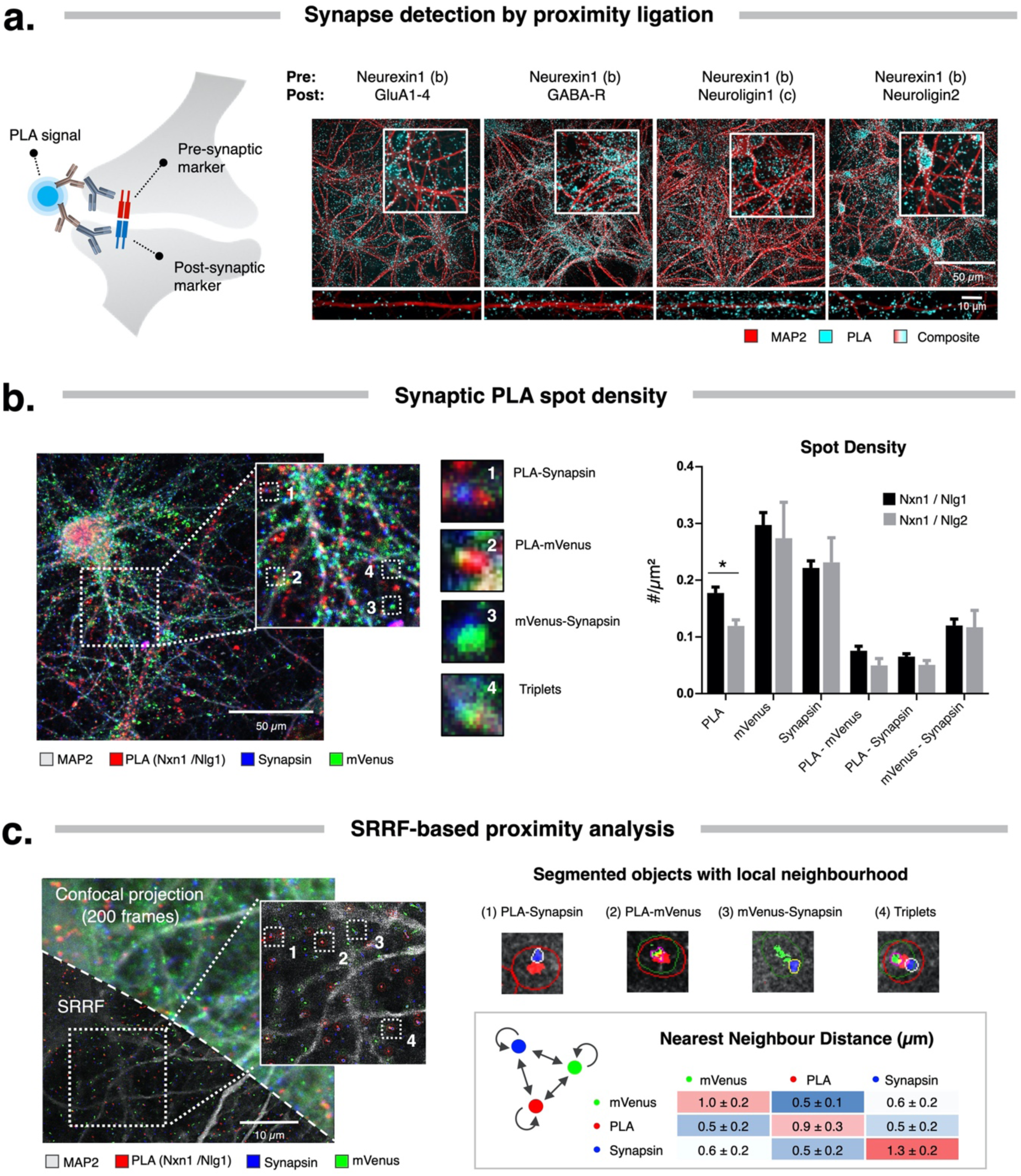
Proximity Ligation Assay for the detection and quantification of synapses. **a.** Diagram illustrating the principle of synaptic PLA (left) and representative images of synaptic PLA signal with the described combination of antibodies used in 14 DIV cortical neurons (right); **b.** Composite image of 28 DIV ENABLED cultures with synaptic PLA for Neurexin-Neuroligin1 and counterstained with MAP2 and Synapsin (triple stained), acquired with standard confocal microscopy. Insets show different types of combinations between synaptic PLA spots, mVenus and Synapsin (center). The adjoined spot density quantification reveals differences between individual markers, pairwise combinations and the PLA partners used (right); **c.** Maximum intensity projection from a single confocal plane of triple stained cortical culture imaged 200 times (top half), overlaid with its superresolution result after application of SRRF (bottom half), along with magnified insets. Nearest neighbor distances were calculated between all spots of two markers, and the mean ± standard deviation is displayed per pairwise combination.

To maximize the efficiency of the PLA reaction, we selected within our panel of antibodies, those that would bind markers that directly interact. We selected Neurexin 1 (b) as the most suitable pan-synaptic marker since it has been proven that it interacts at the molecular level with the extracellular epitopes of 4 postsynaptic proteins with available antibodies, namely: anti-GluA1-4, anti-GABA-R, anti-Neuroligin1 (c), and anti-Neuroligin2 (37–39). Indeed, all 4 combinations yielded characteristic punctate PLA pattern at the expected location: inhibitory synapse markers (GABA-R and Neuroligin2) preferentially located near neuronal somas, whereas excitatory PLA signals (GluA1-4 and Neuroligin1 (c)) were found evenly distributed along the neurites (**Fig. 4a**). ACF plots from synaptic PLA displayed a large range (between 0.82 and 0.93) and small FWHM (ranging between 7.5 to 10.5px) which would set the quality of the staining as one of the best performing compared to single staining antibodies (**Suppl. Fig. 11c**) and outperforming all combinations for double staining.

### Synaptic counterstaining and superresolution reveals synaptic PLA signal offset

We next investigated the subcellular location of synaptic PLA signal for the most optimal PLA pairs Neurexin/Neuroligin1 and Neurexin/Neuroligin2. To this end, we used ENABLED neuronal cultures at 28 DIV and counterstained for the pan-synaptic marker Synapsin (**Fig. 4c**). Synaptic PLA density measurements were significantly higher for Neurexin/Neuroligin1 than Neurexin/Neuroligin2 (0.18 ± 0.02 vs 0.12 ± 0.01 spots/µm^2^; two-way ANOVA, post-hoc Sidak’s test p-value = 0.036). Importantly, the densities of synaptic PLA spots, and synaptic PLA/synapsin, synaptic PLA/mVenus pairs are within the ranges obtained previously by double immunofluorescence colocalisation (0.05 - 0.2 spots/µm^2^) (**Suppl. Fig. 12d**). As mVenus signal is coupled to PSD95, resident in excitatory synapses, we focused on spots densities and overlap of the synaptic PLA for the excitatory combination Neurexin/Neuroligin1. The OC of synaptic PLA signals with mVenus and synapsin was 46%±6 and 39%±2, respectively, indicating that less than half of the PLA signals colocalized with one of these markers. This was lower than the OC between mVenus and synapsin (61% ± 6) (**Suppl. Fig. 12a**). However, when putting the images under scrutiny, we noticed that many of the PLA signals were juxtaposed to the other signals (**Fig. 4b**). To corroborate this, we performed a nearest neighbor correlation analysis between the distributions of detected objects of each marker pair (PLA/mVenus, PLA/synapsin, mVenus/synapsin). In brief, we calculated the probability that the nearest neighbor of a given spot is of a different marker type versus the probability that it could be either of both markers (random distribution). Thus, if both markers were completely independent, the resulting nearest neighbor ratio value (nncorr) would be 1, whereas a completely correlated set would have a ratio value close to 0 (Schematic at **Supl. Fig. 12b**). We found that the nncorr measures for synaptic PLA combinations ranged between 0.05 ± 0.001 and 0.19 ± 0.03, suggesting strong dependence with the other markers, as also confirmed by positive (Homer-Bassoon) and negative (vGAT – PSD95) controls (**Suppl. Fig. 12b**).

To obtain a more resolved view on the actual distances between the PLA signals and the other markers, we employed SRRF (**Fig. 4c**). We found that 95% of the synaptic PLA signals were closer than 230 nm to Synapsin spots (**Fig. 4c, Suppl. Fig 13a**). The average number of Synapsin spots associated with every synaptic PLA spot was 0.95±0.1, which suggests that synaptic PLA spots are in a 1:1 ratio with synapses. When quantifying the spot density in SRRF images, we found that the density of mVenus and Synapsin spots was 0.45 ± 0.13 and 1.08 ± 0.2 spots/µm^2^ respectively, whereas that of synaptic PLA (or PLA with other marker combinations) was ≤ 0.25 spots/µm^2^, similar to IF measurements (**Suppl. Fig. 13b**). Taking advantage of the increased resolution in SRRF images, we measured the average distance between the nearest neighbors from pairs of all marker types. This confirmed that PLA spots were always found in closer proximity to mVenus (0.49 µm) or Synapsin (0.54µm) spots than to other PLA spots (**Fig. 4c**). Despite our restriction to optical sections of 0.4 µm, triplets of synaptic PLA, mVenus and Synapsin were frequently encountered with a density of 0.13 spots/µm^2^ and constituting 46% ± 12 of total synaptic PLA particles (**Suppl. Fig. 13a**). Thus, the density of synaptic PLA spots aligns well with that of best-performing transsynaptic marker combinations, although they occupy a spatial location that is slightly offset from the actual synapses.

### Synaptic PLA increases sensitivity for detecting synapse density changes

As we have previously shown synapse density to scale with culture time (Verschuuren et al., 2019), we decided to further validate synaptic PLA on a set of cultures of increasing maturity (DIV 7, 14, 21). As expected, we found an increase in synapse density (**Fig. 5a**). However, the increase between DIV 7 and DIV 21 as measured by synaptic PLA was ∼2 fold higher than the best co-localization approach (Bassoon/Homer). We next tested whether synaptic PLA could also pick up changes in synapse density as evoked by pathological conditions. We previously found that overexpression of hTau-P301L reduces synapse density from DIV 14 onwards (16). To assess the sensitivity of this assay, we measured synaptic density at 21 DIV after applying increasing multiplicities of infection (300, 400 and 500 MOI) of the hTau-P301L construct. We found a statistically significant decrease of synaptic PLA spot density in hTau-P301L neurons as compared to control cultures (from 0.20 to 0.18, 0.17 and 0.12 spots/µm^2^ for 300, 400 and 500 MOI respectively), which was much more prominent than that of the original colocalization approach (Bassoon-Homer), which only yielded a significant decrease starting for MOI400 (**Fig. 5b**). In light of this, we next assessed whether the synaptic PLA reaction could differentiate between groups where the colocalization approach was not yet sensitive enough. When testing the lowest hTau-P301L construct concentration (300MOI) at an earlier timepoint (DIV14), a statistically significant difference in synaptic density could be detected with synaptic PLA (two-way ANOVA main effect of PLA vs IF (1, 36) = 15.52, p = .003, Sidak’s post-hoc test synaptic PLA CTR vs P301l p<0.001) (**Fig. 5b**). Thus, these results suggest that synaptic PLA allows detecting changes in synapse density more sensitively than the colocalization approach.

**Figure 5.**
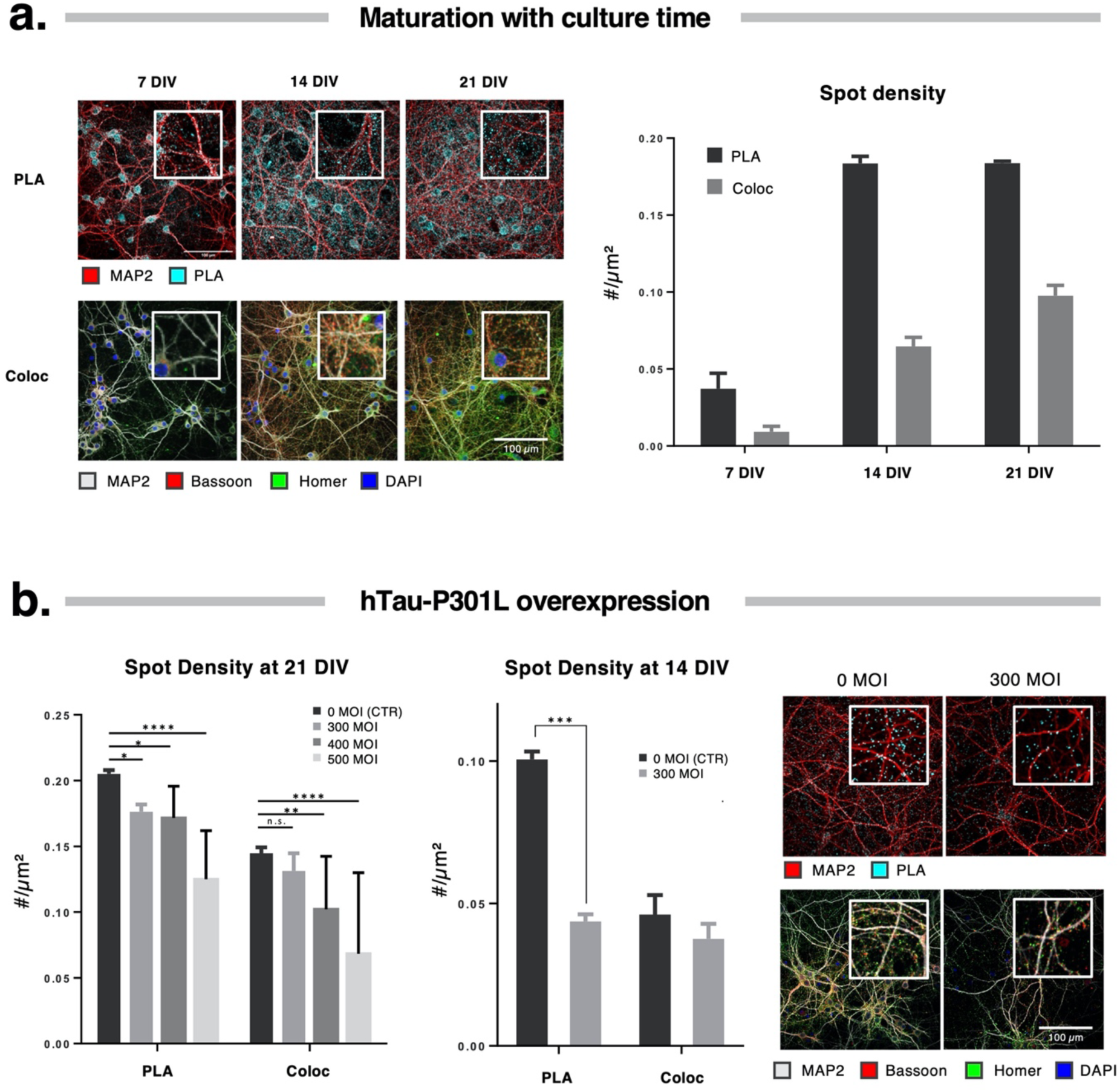
Synaptic PLA detects changes in synapse density in healthy and pathological conditions. **a.** Representative images (left) and quantification of spot density in cortical neuronal cultures at DIV 7, 14 and 21, after synaptic PLA (Neurexin/Neuroligin1) or colocalization (Bassoon/Homer); **b.** Quantification of synapse density after infection with a hTau.P301L AAV vector. Infection with different multiplicities of infection (300, 400 and 500 MOI), reveal a concentration-dependent effect at DIV 21, that is already significant for the lowest MOI when measured with PLA; **c.** Representative images (center) and quantification of neuronal cultures treated with hTau.P301L at 300 MOI revealing a significant effect synapse density at 14 DIV. * = p-value < 0.05; **= p-value < 0.01; *** = p-value < 0.001, **** = p-value < 0.0001, two-way ANOVA post-hoc Sidak’s test.

## Discussion

A wealth of studies has used immunofluorescence to quantify synapse density changes in primary neuronal cultures (6, 8–16, 31, 40). Yet, very few have truly documented the specificity of the utilized antibodies in detail. To gain more certainty about labeling performance, and provide a reference for the field, we have now explored a variety of benchmarking methods to evaluate the specificity of a set of antibodies raised against synaptic proteins. Although extensively used (42), we found that western blotting is not useful for routine antibody validation since neither the presence of additional bands (alternative splice forms) nor the absence of a specific band (epitope masking) can truly predict non-specificity. Therefore, we first introduced a segmentation-independent method to assess the staining performance of individual antibodies in immunofluorescence images. This revealed a large variability between antibodies, which partially depended on the functional class. Synaptic vesicle markers yielded consistently better results in ACF analysis than neurotransmitter receptors or adhesion proteins, likely because the local protein abundance is higher for the former. We further found that the absolute spot density was highly variable even for antibodies that target the same marker. This may be caused by non-specific labeling, low SBR complicating thresholding, and the presence of spurious signals such as markers being trafficked in the neurites or extrasynaptic neurotransmitter receptors. These caveats do not necessarily preclude relative quantification in a setting where experimental conditions are compared to an internal control (8–10), but they may significantly affect sensitivity and render the assay susceptible to confounding factors such as neurotoxicity. By quantifying the association with the postsynaptic compartment, we discovered that Homer outperformed both PSD95 antibodies, both in terms of overlap with the genetic marker PSD95-mVenus and with the presence in dendritic spines. Surprisingly, and in contrast with other reports, we could not recapitulate equally good results with PSD95 antibodies (43). This may be due to the different species (rat vs mouse primary hippocampal cultures) or cultivation protocol (Lagache et al. used astrocyte-conditioned medium). Another explanation may be that Homer, as polyclonal antibody, might be more resistant to epitope masking than PSD95 monoclonals. As exponent, the GluA1-4 antibody showed remarkable low correlation with the postsynaptic compartment, suggesting that many AMPA-R clusters reside outside of the synapse, potentially in reserve pools awaiting synaptic potentiation (44).

We further found that the selectivity for mature synapses could be enhanced by considering the apparent colocalization between a pre- and postsynaptic marker, while maintaining high-content screening compatibility. The positive results from ACF and downstream analyses for Bassoon and Homer antibodies were further substantiated by the high colocalization metrics (CCF amplitude > 0.45, OC > 30%) when used in combination. This makes the latter combination preferred for global assessment of structural connectivity.

While colocalization helps focusing on the relevant fraction of immunolabeled samples, it is subject to bias and suffers from a limited dynamic range. Therefore, we explored whether PLA could serve as a more sensitive alternative. Whereas the PLA technique has been used as a synaptic detection method for tagged proteins, and to localize synaptic proteins to the pre- or post-synaptic compartment (45–47), our work significantly expands the application radius by applying it to transsynaptic connections, rendering it applicable to the detection of mature synapses. Our results demonstrate that synaptic PLA might have additional advantages. First, it allows for much simpler signal detection, as also proven by ACF analysis. Importantly, we found that on average each synaptic PLA spot had ∼1 Synapsin spot, which supports the case that synaptic PLA is a reliable proxy for synapse estimates. Second, using one single marker for synaptic connections frees one channel for additional staining. Finally, our results of synapse quantification in cultures through stages of maturation and pathological perturbations, strongly endorse the potential of synaptic PLA being more sensitive and thus capable of detecting subtler changes in synapse number. This is especially relevant for dense cultures, in which crowding may complicate signal detection. On the downside, one has to consider the increased cost and fast degradation of the PLA signal, which, as yet, may limit its applicability in a high-throughput setting. The offset we encountered between PLA signals and antibody labeling, is most likely due to the amplification protocol, but also should be kept in mind. Another potential issue is the detection of PLA signal in controls that were initially considered to be negative (such as VGAT and Neuroligin1). However, it is becoming clearer that the histological definition of inhibitory vs. excitatory synapses is not clear-cut. For example, in cortical culture, electron microscopy of dendritic spines, once thought to be exclusive of excitatory synapses, also contain inhibitory contacts (32). Additionally, Neuroligin1 and Neuroligin2 have been shown to colocalize with both vGAT and vGlut1, with only enrichment of each isoform on excitatory and inhibitory synapses respectively, but by no means an exclusive colocalization (48).

To conclude, we have extensively validated synapse antibodies in primary cortical and hippocampal culture and demonstrated that synaptic PLA can be more sensitive than synapse detection by marker colocalization. Our work therefore adds a valuable new angle to synapse-oriented *in vitro* screening assays.

## Author contributions

PV, GG, RN, PL and WDV have conceived the work and performed data analysis. PV, GG and BA have performed the wet lab work. MV has developed the synapse quantification script. PV, GG and WDV have drafted the manuscript. All authors have critically revised the manuscript and approved it for publication.

## Acknowledgements

The authors would like to acknowledge Sofie Thys for technical assistance during the preparation of primary neuronal cultures and Shraddha Prasain for Western blotting.

