## Supplementary Information for "Systematic quantification of synapses in primary neuronal culture"

* equal contribution

**Supplementary Figures**

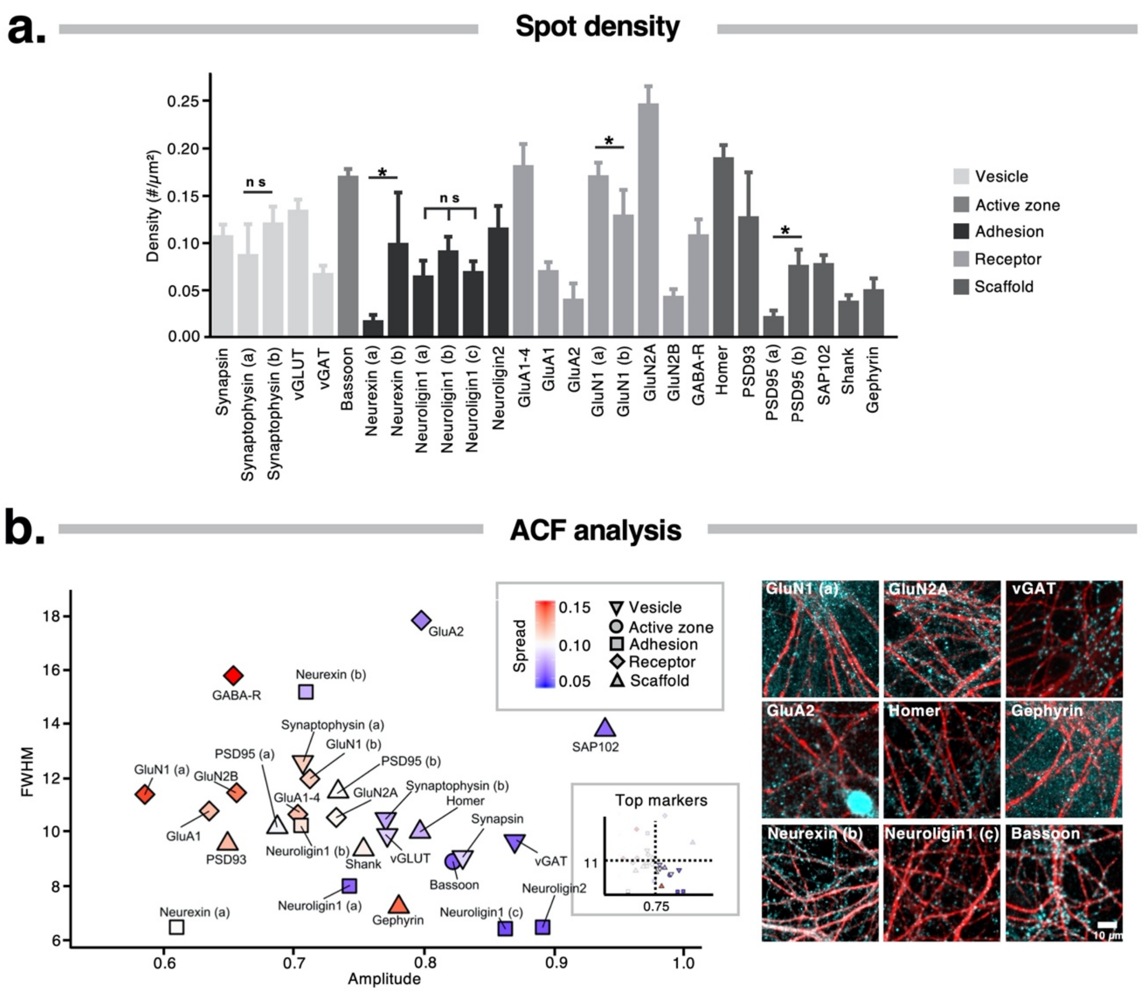


**Supplementary Figure 1.** Synapse marker evaluation in 14 DIV primary hippocampal cultures. **a.** Quantification of the spot density by manual thresholding reveals variable results (*p<0.05, one-way ANOVA post-hoc Sidak's multiple comparisons test); **b.** Synapse antibody evaluation in 14 DIV primary hippocampal cultures. A scatter plot of ACF-derived parameters (and inset with gating on optimal parameter conditions) along with representative images are shown.

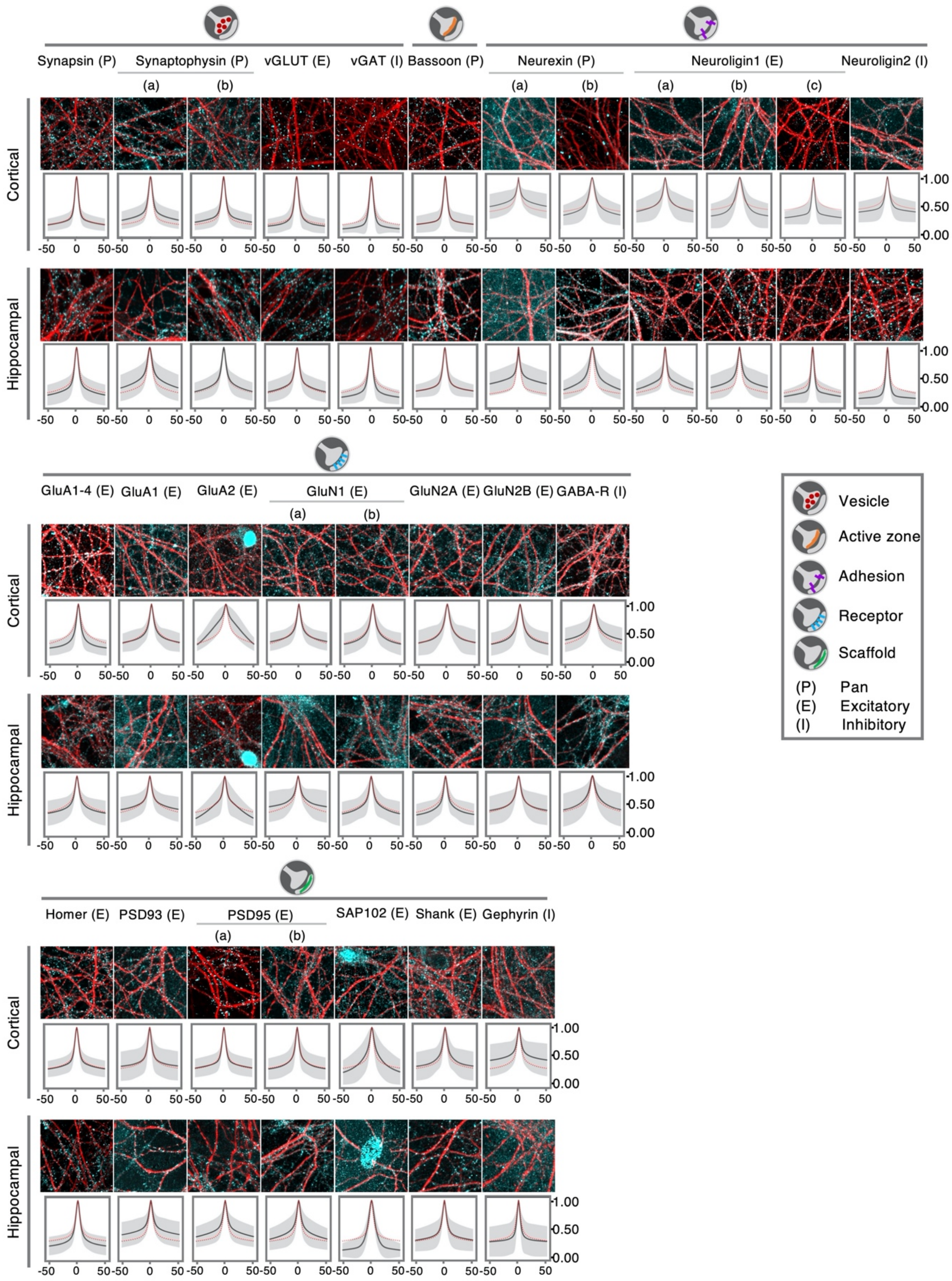


**Supplementary Figure 2.** Labeling specificity of synapse marker antibodies. IF results of all tested antibodies on 14 DIV primary cortical and hippocampal cultures are shown, subdivided in five different functional classes (as per **Fig. 1a**). A representative image of MAP2 (red) with the synapse marker (cyan - contrast optimized) is shown along with the auto-correlation function (ACF) of the image dataset, which served for extracting amplitude, spread and FWHM in **Fig. 1**. The average ACF of the functional class is superimposed in red.

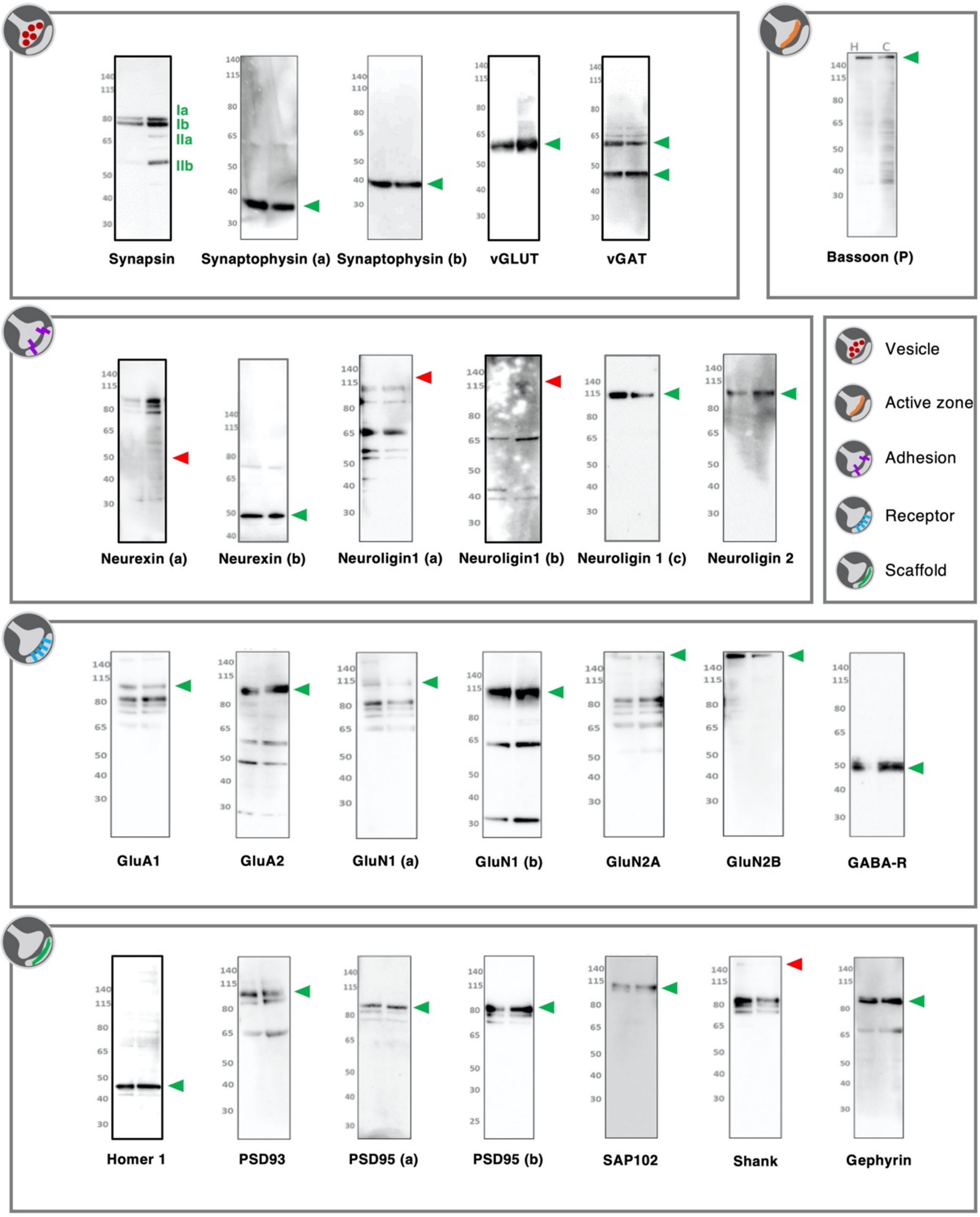


**Supplementary Figure 3.** Antibody validation via western blot. Antibodies were tested on lysates of 14 DIV primary hippocampal (left lane) and cortical (right lane) cultures. An arrowhead indicates the height of the predicted molecular weight of each marker and is shown in green or red when the predicted band is present or absent, respectively.

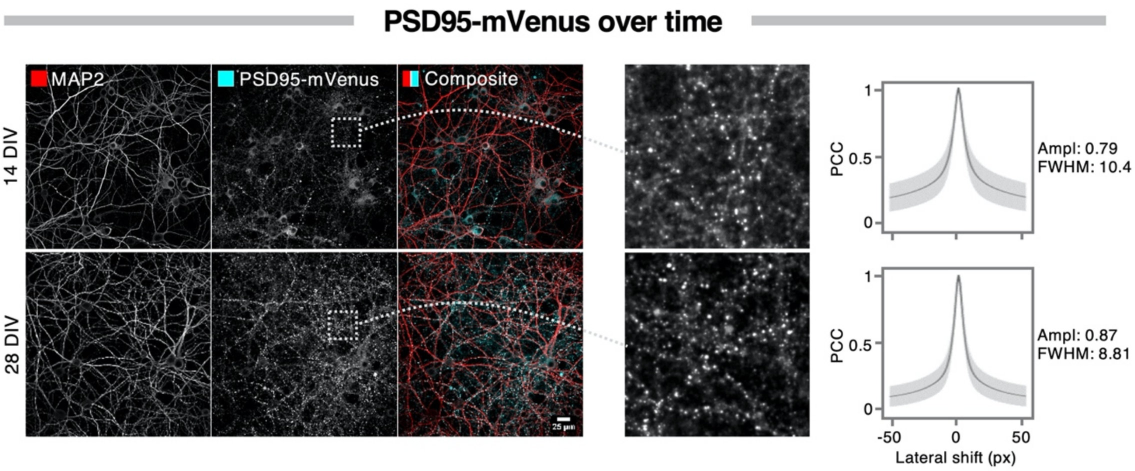


**Supplementary Figure 4.** PSD95-mVenus signal evolves between 14 and 28 DIV. Representative images of 14 and 28 DIV cortical cultures. The contrast-matched images reveal a marked increase in the overall intensity of the PSD95-mVenus signal between 14 and 28 DIV. The insets, shown with optimized contrast settings per DIV, show a larger difference between back- and foreground for 28 as compared to 14 DIV, leading to an ACF with higher amplitude. Note that the ACF relies on the Pearson's Correlation Coefficient (PCC) which is independent of the average intensity.

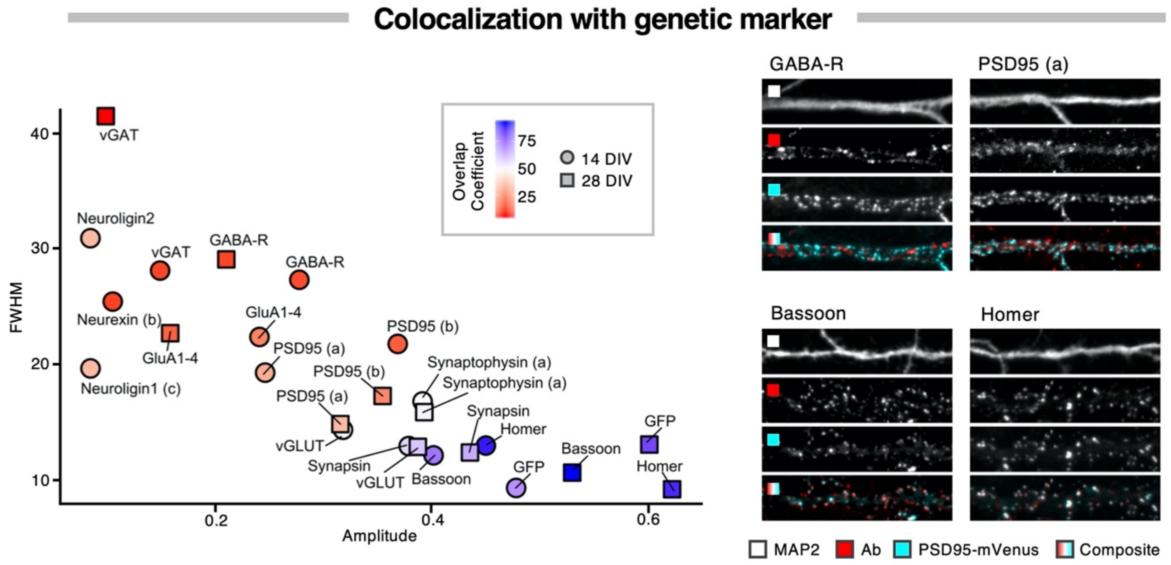


**Supplementary Figure 5.** Colocalization of synapse marker antibodies with PSD95-mVenus in primary hippocampal cultures. The colocalization is shown as a scatter plot of the CCF-derived parameters amplitude (reporting on colocalization) and FWHM (reporting on combined size), and colored by the overlap coefficient (OC), defined as the percentage of PSD95-mVenus spots that have an overlapping antibody spot. Images are from 28 DIV hippocampal cultures.

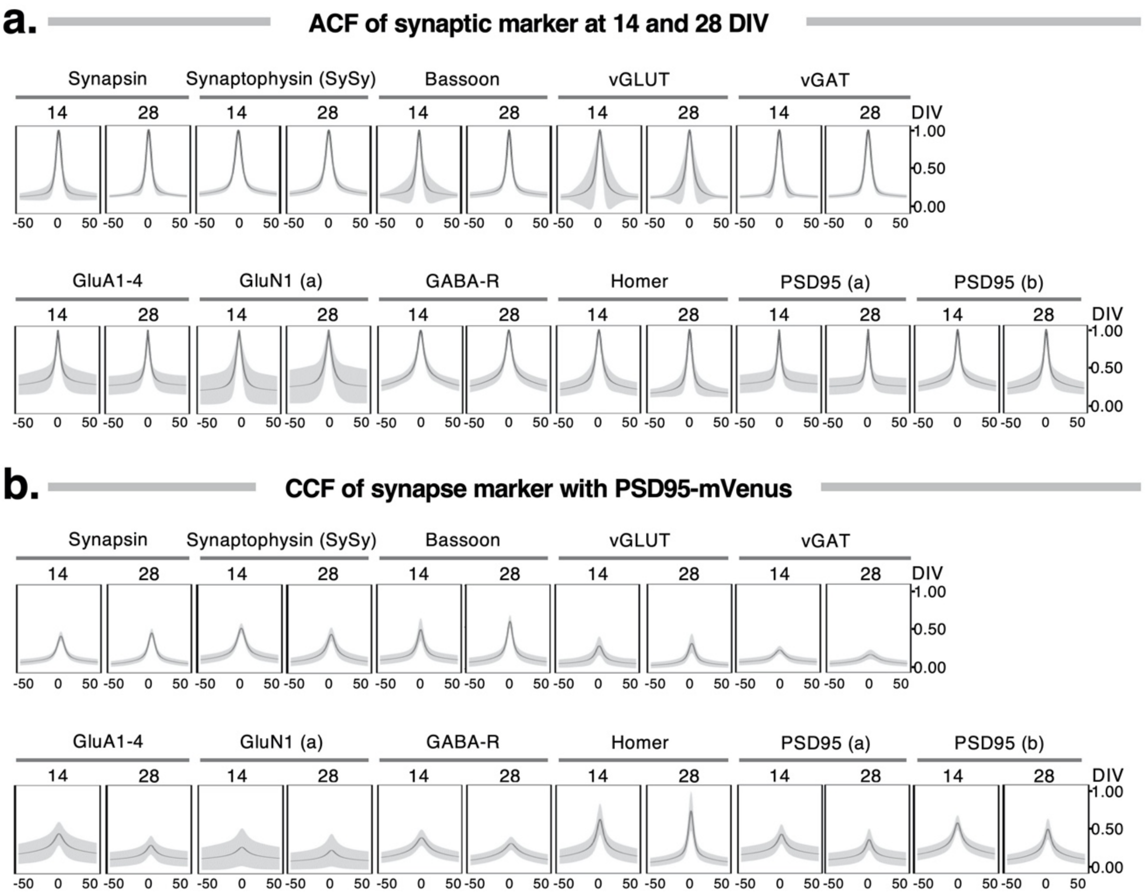


**Supplementary Figure 6.** Auto- and cross-correlation functions of PSD95-mVenus and synapse markers in cortical cultures.

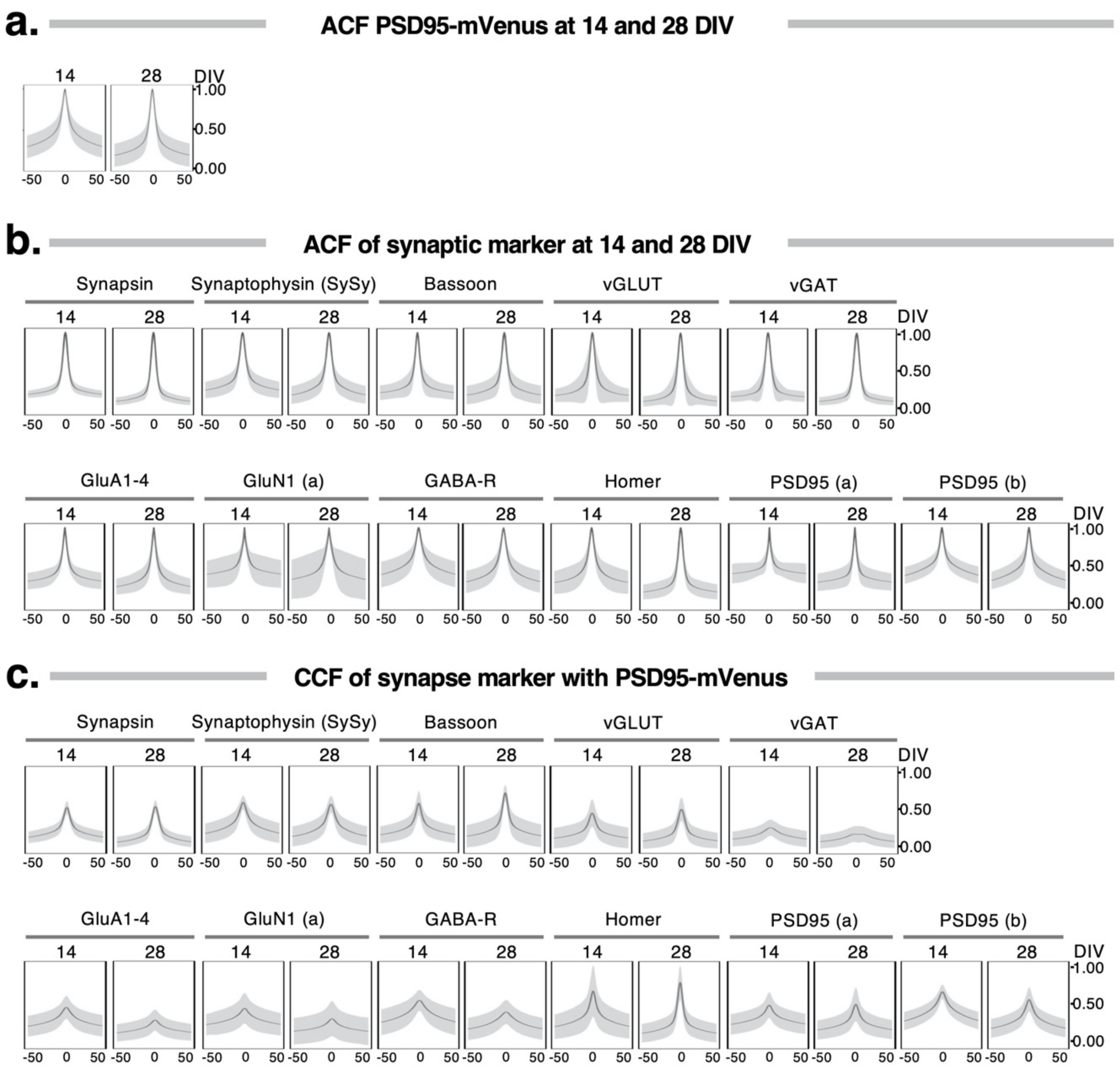


**Supplementary Figure 7.** Auto- and cross-correlation functions of PSD95-mVenus and synapse markers in hippocampal cultures.

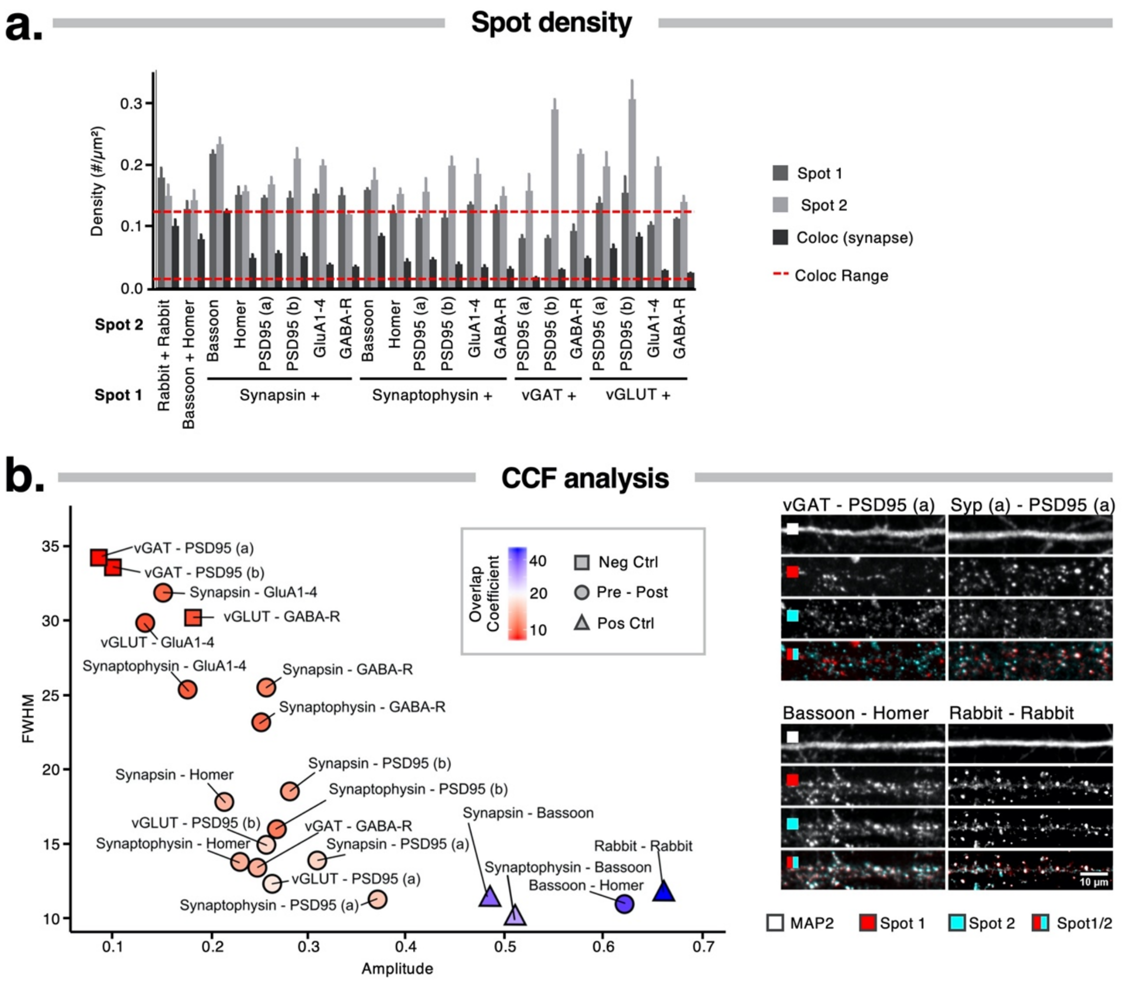


**Supplementary Figure 8.** Double labeling of synapse markers in 14 DIV hippocampal cultures **a.** Quantification of spot and synapse density in 14 DIV cortical cultures shows the limited fraction of colocalized signals; **b.** Representative images and a scatter plot of CCF parameters reporting on the colocalization of marker pairs in 14 DIV hippocampal neurons. The color code indicates the overlap coefficient, defined as the percentage of spots that reside in synapses. Pairs of inhibitory with excitatory synapse markers were considered as negative controls while 2 pan-presynaptic markers and primary antibodies raised in the same species were used as positive controls.

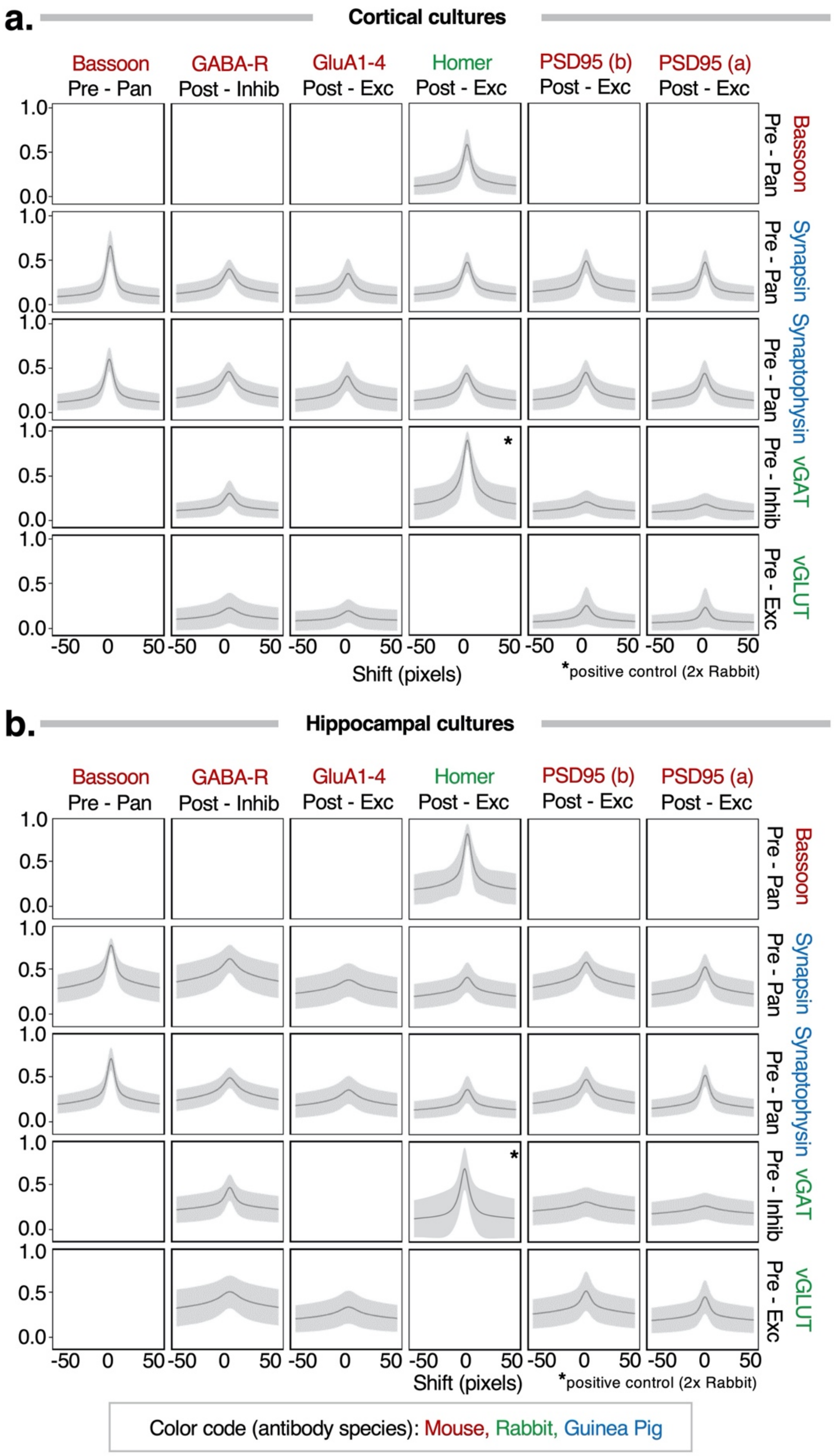


**Supplementary Figure 9.** Cross-correlation functions of double stainings in cortical and hippocampal cultures. CCFs showing the colocalization (amplitude) and combined length (FWHM) of synapse marker combinations in 14 DIV cortical and hippocampal cultures.

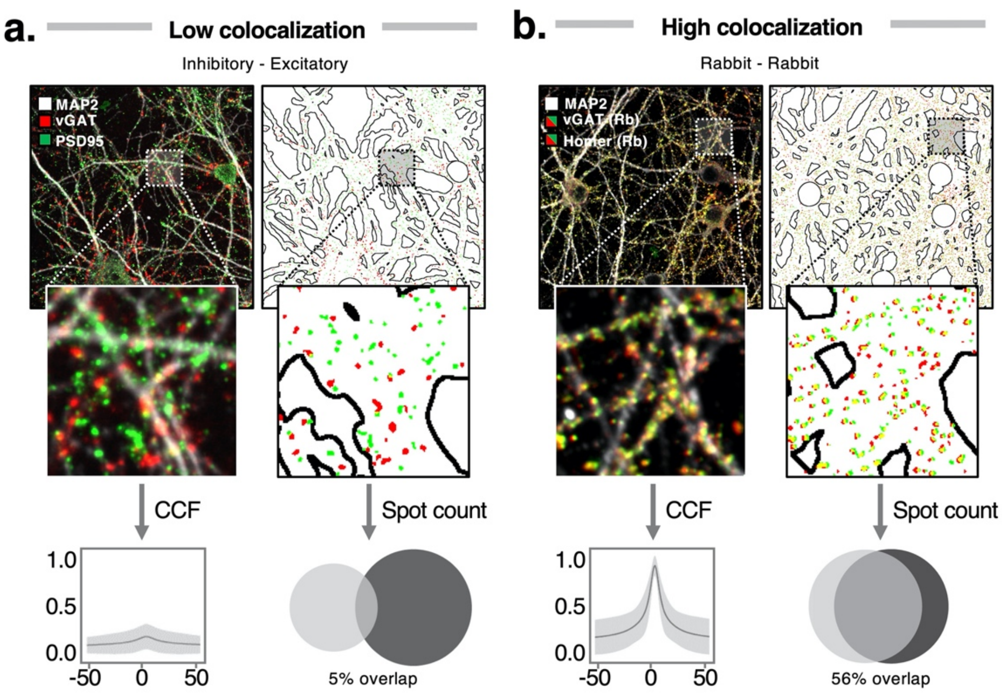


**Supplemental figure 10.** Spot quantification of double stainings with low and high colocalization. The combination of an inhibitory presynaptic (vGAT) with excitatory postsynaptic marker (PSD95) yields a compact CCF and low OC after segmentation. In contrast, a positive control in which 2 rabbit primary antibodies are used, yields a CCF that approximates 1 at $\Delta x = 0$ (nearly perfect colocalization), while the OC is still only 56%, displaying the better sensitivity of the unbiased CCF approach.

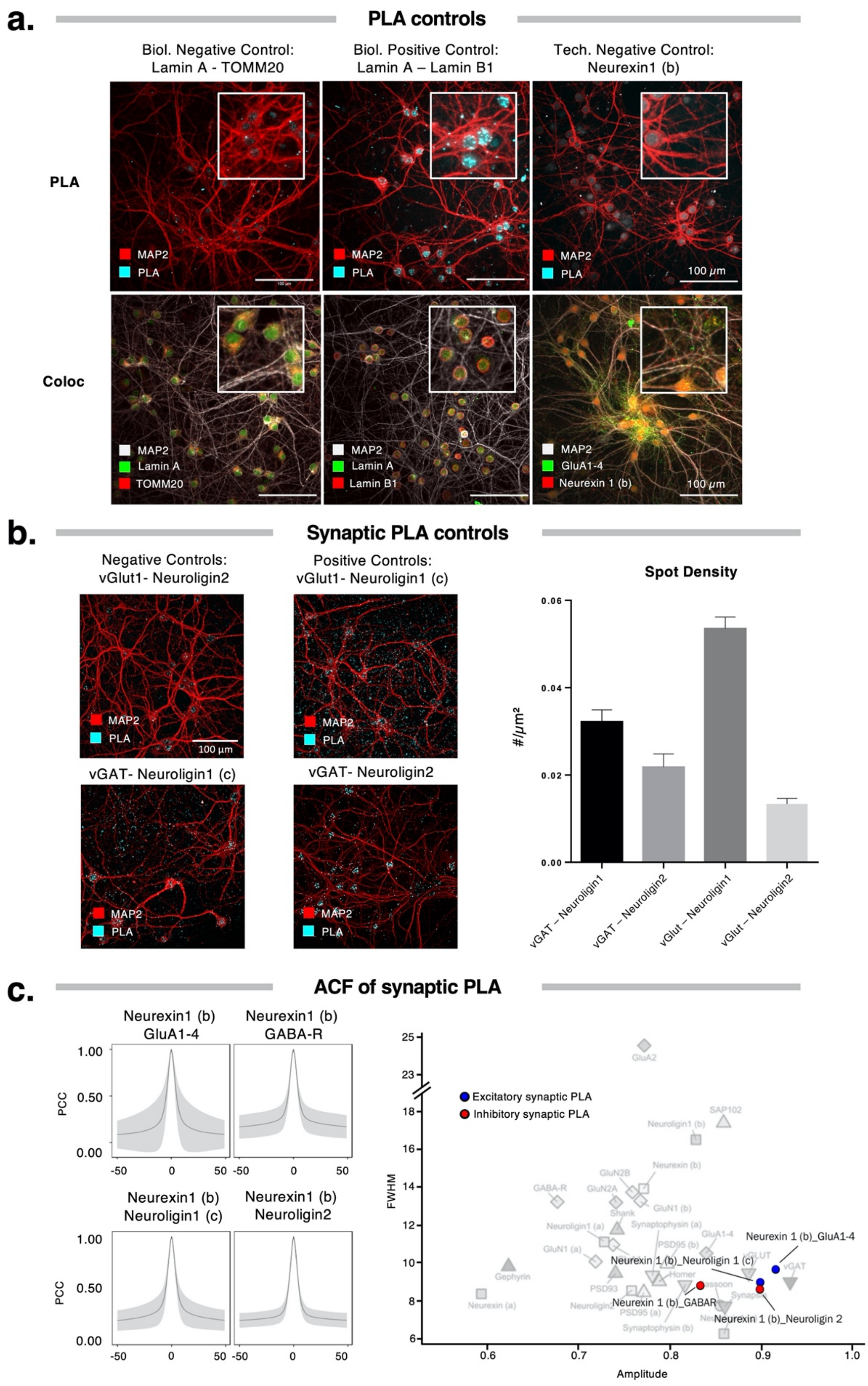


**Supplementary Figure 11.** PLA controls and characteristics. **a.** Positive and negative technical and biological controls show the specificity of PLA in neurons, when comparing the amount of signal and the location of known interacting nuclear proteins (Lamin A / Lamin B1) with proximal but out-of-PLA-reach markers (Lamin A / TOMM20) as well as the technical control of the omission of a singular primary antibody; **b.** Synaptic PLA with permutations of inhibitory and excitatory markers show that congruent combinations produce increased numbers of PLA spots; **c.** ACF plots for PLA signal from specified markers (left); and scatterplot location of synaptic PLA marker combinations as compared with single-staining ACF properties (right, cfr. also **Fig. 1**).

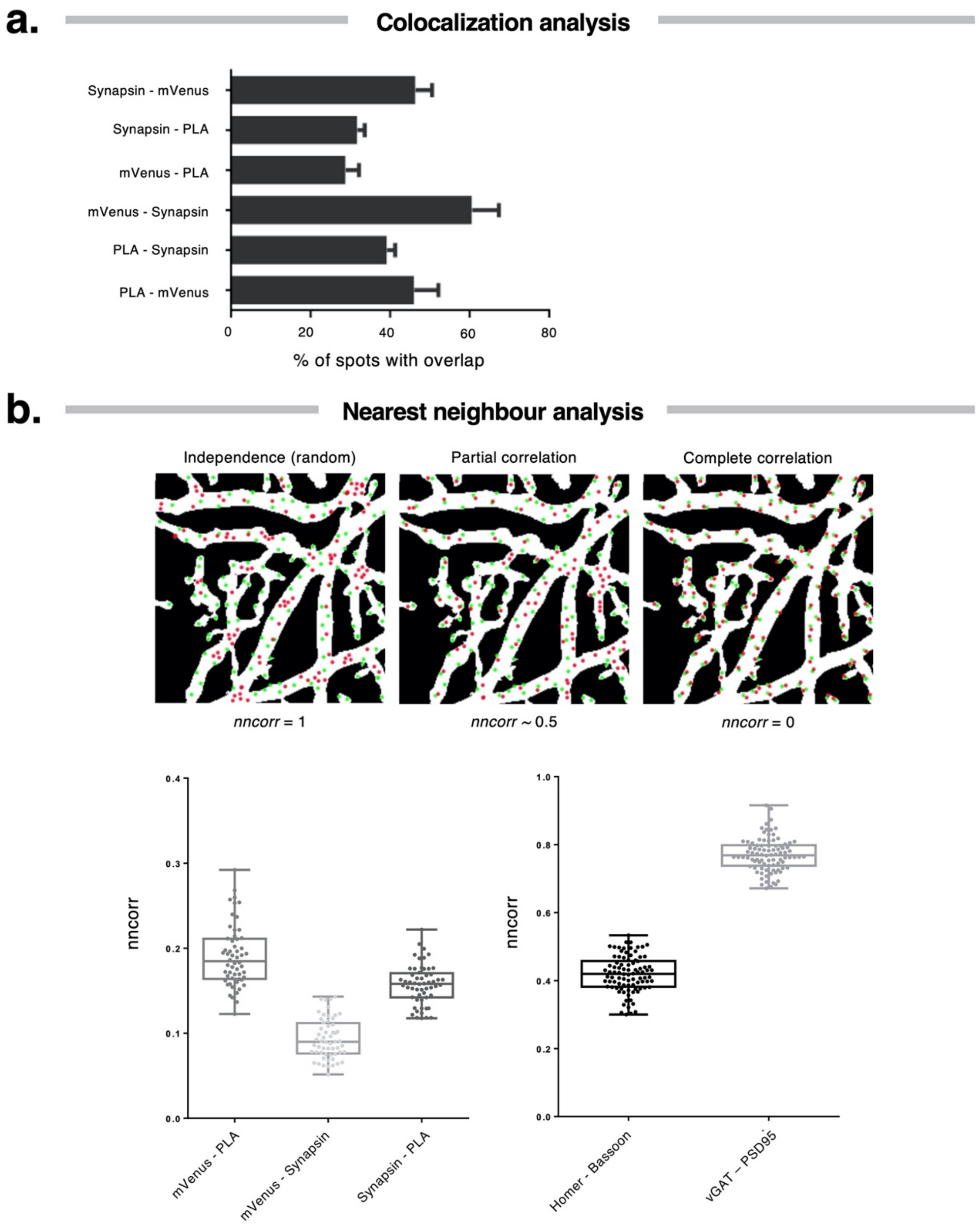


**Supplementary Figure 12.** Analyses of PLA triple staining (synaptic PLA-Neurexin/Neuroligin1, mVenus-PSD95 and synapsin) of cortical 28DIV cultures. **a.** Quantification graph of overlap percentage for the specified pairs of spots derived from the segmentation of confocal images. Values are the percentage of each pair against the total number by marker (*e.g.,* PLA – mVenus is the percentage of PLA spots that have an overlapping mVenus signal); **b.** Nearest neighbor correlation schematic and quantification. *Nncorr* measures the relationship between two types of spots by calculating the proportion of spots that have a nearest neighbor of the same type, and normalizing by what would be expected from an independent (random) distribution; thus values close to 1 suggest independence between the sets of spots, whereas a value closer to 0 is expected if spots are correlated. PLA triple staining pairs (mVenus-PLA, PLA-synapsin and mVenus-synapsin), present lower *nncorr* values implying that spots are correlated. This was also validated by comparing positive (Homer/Basoon) and negative (VGAT/PSD95) control IF combinations. Note that in the latter case a different microscopy setup was used, leading to allover higher values.

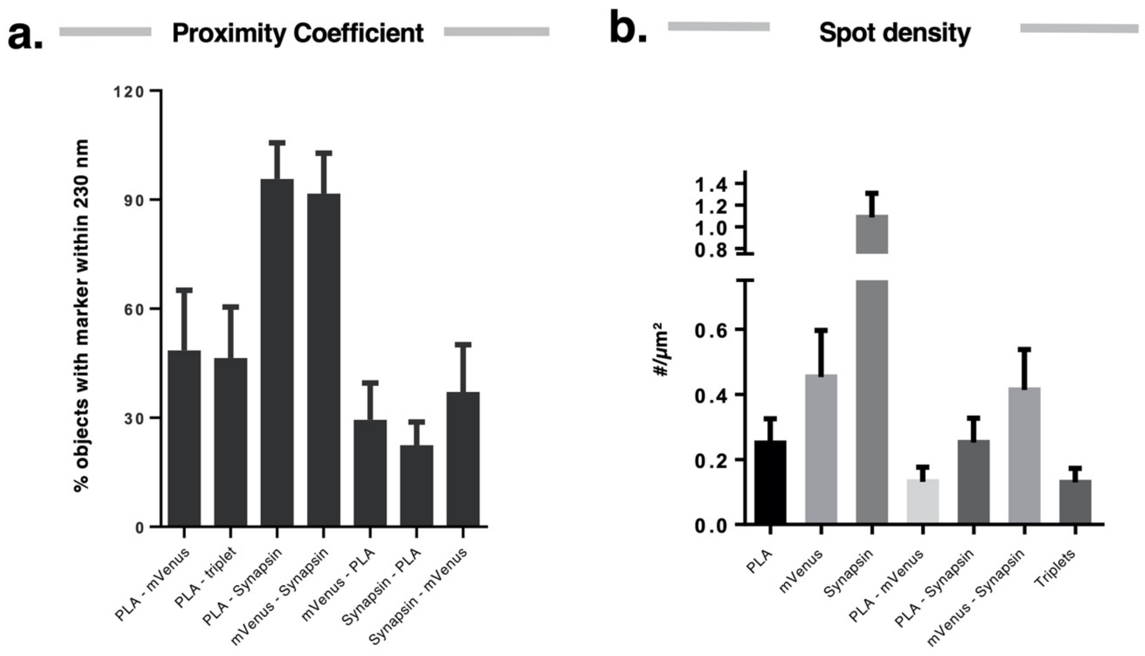


**Supplementary Figure 13.** Quantification of spots from SRRF images with triple staining (synaptic PLA, Synapsin and mVenus-PSD95). **a**. Quantification of proximity for the specified spots. Spot objects were identified in SRRF images of a single confocal plane of cortical 28 DIV cultures and were then expanded by 10 pixels (~230nm) to identify juxtaposed spots. Values represent the percentage of spots with juxtaposition of another marker spot versus the total number of spots (*e.g.,* PLA – mVenus is the percentage of PLA spots that have an mVenus spot within 230 nm); **b.** Spot density for each staining, pairwise combinations and triplets, derived from SRRF images on PLA triple stained cortical 28 DIV cultures.

**Table 1.** Primary and Secondary antibodies employed in this study (antibodies used in PLA are marked with an asterisk)

**PRIMARY ANTIBODIES**

| Antigen | Pre/Post | Type | Localization | Epitope | Species/Clonality | Clone | Company | Catalog | Lot | IF conc. | WB conc. |
|---|---|---|---|---|---|---|---|---|---|---|---|
| Bassoon | Pre | Pan | Active zone | intracellular | Mouse Monoclonal | L124/59 | Neuromab | 73-491 | 463-3DH-50 | 2 µg/ml | 3 µg/ml |
| Neurexin 1β (a) | Pre | Pan | Adhesion | extracellular | Mouse Monoclonal | N170A/1 | Neuromab | 75-216 | 447-1JH-73A | 20 µg/ml | 1 µg/ml |
| Neurexin 1* (b) | Pre | Pan | Adhesion | Intracellular | Rabbit Polyclonal | NA | Abcam | Ab222806 | GR3192323-13 | 40 µg/ml | 1 µg/ml |
| Synapsin 1/2 | Pre | Pan | Vesicular | intracellular | Guinea Pig Polyclonal | NA | SySy | 106004 | 106004/1-21 | 2 µg/ml | 0.5 µg/ml |
| Synaptophysin 1 (a) | Pre | Pan | Vesicular | intracellular | Guinea Pig Polyclonal | NA | SySy | 101004 | 101004/2-26 | 2 µg/ml | 0.25 µg/ml |
| Synaptophysin 1 (b) | Pre | Pan | Vesicular | intracellular | Rabbit Polyclonal | NA | Abcam | ab14692 | GR200179-2 | 2 µg/ml | 0.65 µg/ml |
| vGAT | Pre | Inhibitory | Vesicular | intracellular | Rabbit Polyclonal | NA | SySy | 131003 | 131003/1-45 | 2 µg/ml | 0.25 µg/ml |
| vGlut1 | Pre | Excitatory | Vesicular | intracellular | Rabbit Polyclonal | NA | SySy | 135303 | 135303/2-63 | 2 µg/ml | 0.25 µg/ml |
| AMPA-R (GluA1) | Post | Excitatory | Receptor | extracellular | Mouse Monoclonal | N355/1 | Neuromab | 75-327 | 455-5JD-89 | 2 µg/ml | 1 µg/ml |
| AMPA-R (GluA1-4)* | Post | Excitatory | Receptor | extracellular | Mouse Monoclonal | 248B7 | SySy | 182411 | 182411/10 | 2 µg/ml | 1 µg/ml |
| AMPA-R (GluA2) | Post | Excitatory | Receptor | intracellular | Mouse Monoclonal | L21/32 | Neuromab | 75-002 | 455-6JD-81C | 2 µg/ml | 1 µg/ml |
| GABA-R (β2/3)* | Post | Inhibitory | Receptor | extracellular | Mouse Monoclonal | 62-3G1 | Neuromab | 75-363 | 455-2JD-15 | 2 µg/ml | 1 µg/ml |
| Gephyrin | Post | Inhibitory | Scaffold | intracellular | Mouse Monoclonal | L106/93 | Neuromab | 75-444 | 455-8JD-47 | 2 µg/ml | 1 µg/ml |
| Homer-1 | Post | Excitatory | Scaffold | intracellular | Rabbit Polyclonal | NA | SySy | 160003 | 160003/31 | 2 µg/ml | 1 µg/ml |
| Neuroligin 1 (a) | Post | Excitatory | Adhesion | intracellular | Mouse Monoclonal | N97A/31 | Neuromab | 75-160 | 443-2KS-20C | 20 µg/ml | 1 µg/ml |
| Neuroligin 1 (b) | Post | Excitatory | Adhesion | extracellular | Rabbit Polyclonal | NA | Almone | ANR-035 | ANR035AN0125 | 8 µg/ml | 1 µg/ml |
| Neuroligin 1* (c) | Post | Excitatory | Adhesion | intracellular | Mouse Monoclonal | A-4 | Santa Cruz | Sc-365110 | B0618 | 4 µg/ml | 1 µg/ml |
| Neuroligin 2* | Post | Inhibitory | Adhesion | intracellular | Mouse Monoclonal | L107/39 | Neuromab | 75-451 | 455-8JD-93 | 20 µg/ml | 1 µg/ml |
| NMDA-R (GluN1) (a) | Post | Excitatory | Receptor | extracellular | Mouse Monoclonal | N308/48 | Neuromab | 75-272 | 455-7JD-90B | 2 µg/ml | 1 µg/ml |
| NMDA-R (GluN1) (b) | Post | Excitatory | Receptor | extracellular | Mouse Monoclonal | M68 | SySy | 114011 | 114011/1-24 | 2 µg/ml | 0.25 µg/ml |
| NMDA-R (GluN2A) | Post | Excitatory | Receptor | extracellular | Mouse Monoclonal | N327/95 | Neuromab | 75-288 | 455-7JD-90B | 2 µg/ml | 1 µg/ml |
| NMDA-R (GluN2B) | Post | Excitatory | Receptor | extracellular | Mouse Monoclonal | N59/20 | Neuromab | 73-097 | 463-1DH-64 | 2 µg/ml | 2 µg/ml |
| PSD93 | Post | Excitatory | Scaffold | intracellular | Mouse Monoclonal | N18/30 | Neuromab | 75-057 | 455-3JD-72 | 2 µg/ml | 1 µg/ml |
| PSD95 (a) | Post | Excitatory | Scaffold | intracellular | Mouse Monoclonal | 7E3-1B8 | ThermoFisher | MA1-046 | SA242215 | 2 µg/ml | 0.5 µg/ml |
| PSD95 (b) | Post | Excitatory | Scaffold | intracellular | Mouse Monoclonal | K28/43 | Neuromab | 75-028 | 455-7JD-22F | 2 µg/ml | 1 µg/ml |
| SAP102 | Post | Excitatory | Scaffold | intracellular | Mouse Monoclonal | N19/2 | Neuromab | 75-058 | 449-1AK-85B | 2 µg/ml | 1 µg/ml |
| Shank 1/2/3 | Post | Excitatory | Scaffold | intracellular | Mouse Monoclonal | N23B/49 | Neuromab | 75-089 | 443-1KS-78B | 2 µg/ml | 1 µg/ml |
| MAP2 | NA | NA | Dendrite | intracellular | Chicken Polyclonal | NA | SySy | 188006 | 188006/1-3 | 2 µg/ml | NA |
| GAPDH | NA | NA | Cytoplasm | intracellular | Mouse Monoclonal | GT239 | GeneTex | GTX627408 | 41323 | NA | 1 µg/ml |
| GFP/mVenus | NA | NA | NA | intracellular | Alpaca Monoclonal | +Atto594 | Chromogen | Gba594 | 81212001AT3 | 1 µg/ml | NA |

**SECONDARY ANTIBODIES**

| Type | Conjugate | Company | Catalog | Lot | Concentration |
|---|---|---|---|---|---|
| Donkey-anti-Guinea Pig | HRP | Jackson Immunoresearch | 706-035-148 | 128221 | 0.1 µg/ml |
| Donkey-anti-Mouse | HRP | Jackson Immunoresearch | 715-035-151 | 125909 | 0.5 µg/ml |
| Donkey-anti-Rabbit | HRP | Jackson Immunoresearch | 711-035-152 | 104907 | 0.2 µg/ml |
| Donkey-anti-Chicken | AlexaFluor647 | Jackson Immunoresearch | 703-605-155 | 134612 | 1 µg/ml |
| Goat-anti-Mouse | AlexaFluorPlus488 | ThermoFisher | A32723 | SG251135 | 2 µg/ml |
| Goat-anti-Rabbit (Fab) | FITC | Jackson Immunoresearch | 111-097-003 | 104632 | 0.5 µg/ml |
| Donkey-anti-Guinea Pig | Cy3 | Jackson Immunoresearch | 706-165-148 | 127715 | 1 µg/ml |
| Donkey-anti-Mouse | Cy3 | Jackson Immunoresearch | 715-165-151 | 125797 | 1 µg/ml |
| Goat-anti-Rabbit (Fab) | Cy3 | Jackson Immunoresearch | 111-167-003 | 78942 | 0.5 µg/ml |
